# PKD autoinhibition in *trans* regulates activation loop autophosphorylation in *cis*

**DOI:** 10.1101/2022.05.05.490744

**Authors:** Ronja Reinhardt, Kai Hirzel, Gisela Link, Stephan A. Eisler, Tanja Hägele, Matthew A.H. Parson, John E. Burke, Angelika Hausser, Thomas A. Leonard

## Abstract

Phosphorylation is a ubiquitous mechanism by which signals are transduced in cells. Protein kinases, enzymes that catalyze the phospho-transfer reaction are, themselves, often regulated by phosphorylation. Paradoxically, however, a substantial fraction of the more than 500 human protein kinases are capable of catalyzing their own activation loop phosphorylation. Commonly, these kinases perform this autophosphorylation reaction in *trans*, whereby transient dimerization leads to the mutual phosphorylation of the activation loop of the opposing protomer. In this study, we demonstrate that Protein Kinase D (PKD) is regulated by the inverse mechanism of dimerization-mediated *trans*-autoinhibition, followed by activation loop autophosphorylation in *cis*. We show that PKD forms a stable face-to-face homodimer that is incapable of either auto- or substrate phosphorylation. Dissociation of this *trans*-autoinhibited dimer results in activation loop autophosphorylation, which occurs exclusively in *cis*. Phosphorylation serves to increase PKD activity and prevent *trans*-autoinhibition, thereby switching PKD on. Our findings not only reveal the mechanism of PKD regulation, but have profound implications for the regulation of many other eukaryotic kinases.

## Introduction

Protein phosphorylation is the most abundant and important post translational modification in cellular signal transduction ^1^. Hence, protein kinases are versatile switches in countless signaling cascades and their dysregulation is a major threat to the fidelity of cell signaling and, eventually, the health of an organism. Protein kinases are, therefore, tightly regulated enzymes that integrate various signaling inputs to drive downstream signaling in the form of substrate phosphorylation. Whilst the active conformation of a kinase domain is conserved amongst the entire class of enzymes, the inactive conformations that are used to suppress their activity in the absence of a physiological stimulus are diverse ^2^. Importantly, the inactive state of a kinase is not just the absence of its active conformation, but well-defined and tightly regulated conformations that rely on a limited repertoire of reoccurring patterns ^3^. As such, the inactive conformation can represent a unique and highly specific target for pharmacological intervention ^4^, while inhibitors that target the highly conserved active conformation are prone to off target effects ^5^.

A major determinant for the activity status of a protein kinase is the trajectory of a flexible loop within the kinase, called the activation loop. While its inactive conformation of which can vary greatly ^3^, active kinases, in contrast, adopt a single, well-defined conformation of their activation loop ^2^. The acquisition of the active conformation is, in many eukaryotic protein kinases, accomplished through phosphorylation of a conserved serine, threonine or tyrosine residue in the activation loop by an upstream kinase. Phosphorylation creates a network of hydrogen bonds and electrostatic interactions that stabilizes the packing of an otherwise labile loop against the kinase domain, which both creates the surface for substrate binding and organizes the catalytic machinery for productive phospho-transfer ^2, 6^.

A special case is kinases that phosphorylate their own activation loop. While more than 18% of protein kinases have been reported to autophosphorylate their activation loop ^7^, the mechanisms by which they acquire autocatalytic activity are still debated. Multiple models have been proposed that cover activation loop autophosphorylation in both *cis* and *trans* ^7, 8^, although the majority of kinases that autophosphorylate are believed to do so in *trans* ^7^. Logic implies that these kinases possess an intrinsic capacity to autophosphorylate in the absence of phosphorylation of their own activation loop. Therefore, the autophosphorylation reaction is necessarily mechanistically distinct from the classical mode of substrate phosphorylation.

One example of a kinase reported to autophosphorylate is Protein Kinase D (PKD). PKD is a family of Ser/Thr kinases of the calcium/calmodulin-dependent kinase (CAMK) family comprising three isoforms, PKD1, 2 and 3, which all share the same domain arrangement (Figure 1A). In epithelial cells, PKD is located at the *trans*-Golgi network (TGN), where its activity is required for the formation of secretory cargo carriers of the TGN to the cell surface (CARTS) ^9, 10^. PKD activation depends on binding of its C1 domains to diacylglycerol (DAG) in the TGN ^11, 12^, which results in its autophosphorylation on S742 and activation ^13, 14^. A secondary, non-canonical, phosphorylation site (S738) in the activation loop of PKD1 has been reported ^13, 15^, though its physiological relevance is not clear. Once activated, PKD regulates vesicular transport by phosphorylating and activating the lipid kinase PI4KB ^16^, among other substrates ^17, 18^. In previous work we have shown that activation loop phosphorylation of PKD is dependent on a ubiquitin-like dimerization domain (ULD) in its N-terminus ^14^. However, the structural mechanism underlying phosphorylation-mediated PKD activation remained elusive.

**Figure 1.**
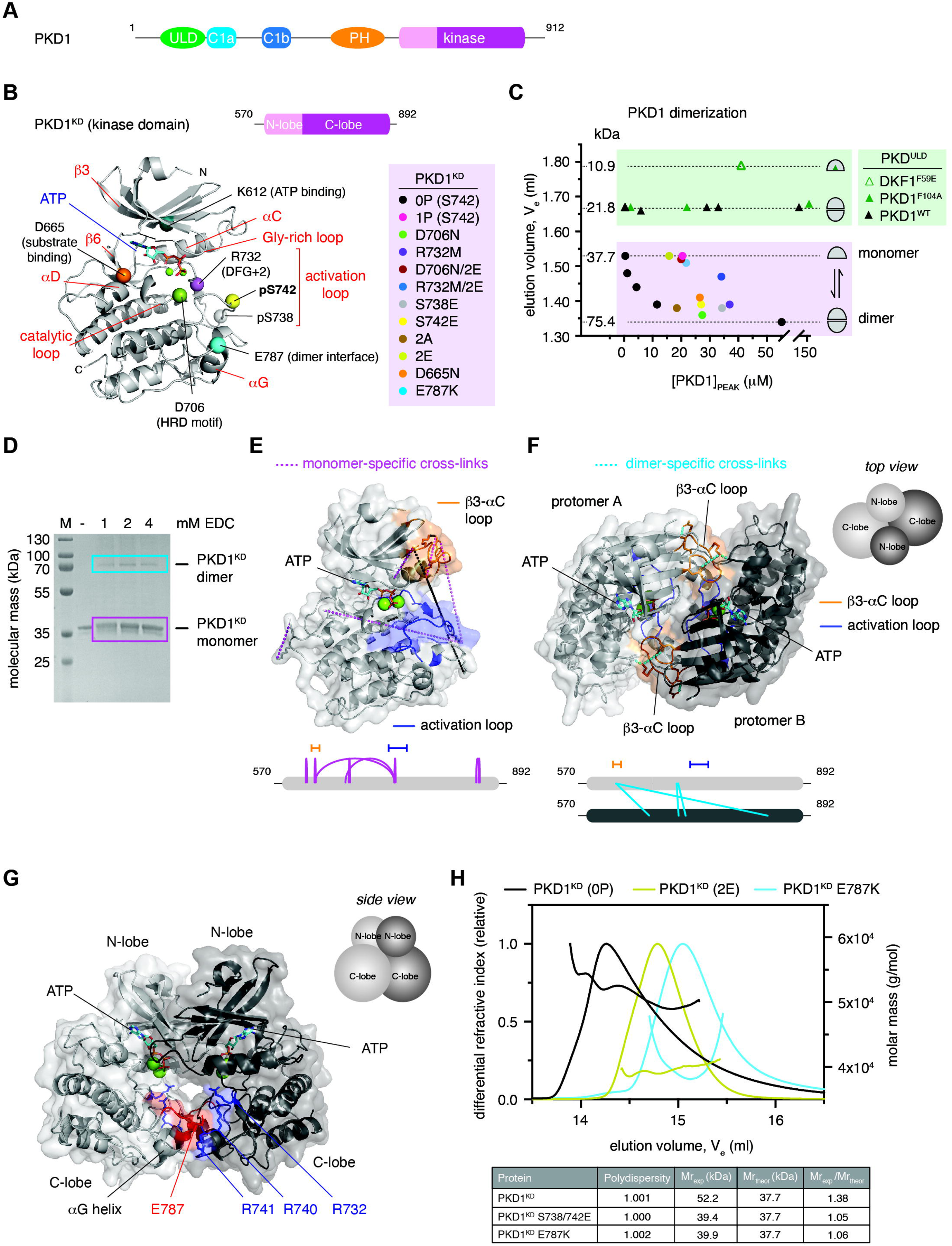
The PKD kinase domain forms a face-to-face dimer. A. Domain architecture of human PKD1. B. PKD variants (color-coded) employed in this study are mapped onto a homology model of the PKD1 kinase domain (PKD1^KD^). C. Analytical size-exclusion chromatography of PKD1^ULD^ and PKD1^KD^ variants. Dashed lines correspond to monomer and dimer, calibrated by multi-angle light scattering to determine absolute molecular mass. D. Cross-linking of PKD1^KD^ with 1-ethyl-3-(3-dimethylaminopropyl)carbodiimide (EDC). Monomer (magenta box) and dimer (cyan box) bands were subjected to mass spectrometry analysis. E. Monomer-specific cross-links mapped onto the Rosetta homology model of PKD1^KD^. Activation loop (blue) and β3-αC loop (orange) are indicated. Below: schematic of EDC cross-links specific to the monomer band (magenta box in D). F. Dimer-specific cross-links mapped onto the cross-link constrained model of the PKD1^KD^ dimer obtained from comparative modeling in Rosetta with C2 symmetry. Below: schematic of EDC cross-links specific to the dimer band (cyan box in D). G. Side view of the experimentally constrained dimer model of PKD1^KD^. The conserved acidic patch at the tip of the αG helix (red) and conserved basic residues in the activation loop (blue) are indicated. H. Size-exclusion chromatography coupled to multi-angle light scattering (SEC-MALS) of wild type (black), S738E/S742E (yellow-green), and E787K (cyan) variants of PKD1^KD^.

In this study, we biochemically dissect the mechanisms of autoinhibition and activation of PKD. We show that the kinase domain of PKD, counterintuitively, forms an inactive dimer. We demonstrate that activation loop auto-phosphorylation is promoted by dissociation of the inactive dimer and that phosphorylation both increases kinase activity and prevents reassociation of the kinase domain. We present biochemical evidence that autophosphorylation and substrate phosphorylation are mechanistically distinct and that activation loop autophosphorylation occurs, not in *trans*, but in *cis*. Finally, we show that dissociation of the kinase domains of inactive PKD dimers and their subsequent activation loop autophosphorylation is essential for constitutive secretion in cells.

## Results

### The PKD kinase domain forms a face-to-face dimer

PKD activation and autophosphorylation in cells is dependent on the formation of a homodimer that is mediated via its N-terminal ULD and C-terminal kinase domain. The unphosphorylated kinase domain of PKD forms a stable dimer in solution, while activation loop phosphorylation prevents its dimerization ^14^, but the mechanism by which phosphorylation activates PKD is, as yet, unknown. To obtain structural insights into the dimerization of the kinase domain of PKD, we built *in silico* models of both the monomeric and dimeric states of the kinase domain, guided by a combination of careful site directed mutagenesis (Figure 1B), oligomeric state determination (Figure 1C), in-solution crosslinking (Figure 1D), and evolutionary sequence conservation.

All recombinant proteins used in this study were purified to homogeneity and subsequently analyzed by intact mass spectrometry to confirm their purity and precise chemical identity (Figure S1-S2). The PKD1 kinase domain construct, henceforth referred to as PKD1^KD^, comprises residues 570-890 and includes, in addition to the canonical kinase domain, short flanking regions with high sequence conservation in vertebrate PKD orthologs. The oligomeric state of each protein was evaluated by analytical size exclusion chromatography (Figure 1C), for which the correlation of elution volumes to molecular masses was calibrated by size exclusion chromatography coupled to multi-angle light scattering (SEC-MALS).

We first used the comparative modelling tool of the Rosetta in silico modeling suite ^19^ to predict the structure of the PKD1 kinase domain based on published crystal structures of the related protein kinases Chk2 (pdb:3i6U), DAPK2 (pdb:2a2a, 2yaa), and phosphorylase kinase (2phk), taking into account the correct geometry and stereochemistry of ATP and Mg^2+^ binding (Figure 1E). Comparison of the resulting model to the AlphaFold prediction (AF-Q15139-F1) ^20^ shows strong agreement, with an overall RMSD of 0.680 Å over all C_α_ atoms (Supplementary Figure S3A).

To provide experimental restraints for *in silico* modelling of the dimeric state we performed in-solution crosslinking with the heterobifunctional, zero-length crosslinker 1-ethyl-3-(3-dimethylaminopropyl)carbodiimide hydrochloride (EDC) on dephosphorylated PKD1^KD^ (Figure 1D). Since thermal stability measurements of the PKD kinase domain indicated the strongest stabilization by ATP (Supplementary Figure S3B), cross-linking was performed under conditions of saturating ATP and MgCl_2_. Mass spectrometric analysis of crosslinked residues in both the monomeric and the dimeric species isolated from the gel (Figure 1D) revealed several major changes upon dimerization (Figure 1E-F, Supplementary Table S1). Crosslinks from the activation loop residue K737 to E668 in the αD helix and E624 in the β3-αC loop were unique to the monomer, while crosslinking of E736 in the activation loop and K622 in the β3-αC loop was reduced in the dimer. Together with an abundance of monomer-specific loop-links that are not observed in the dimer, this indicates a pronounced loss of activation loop flexibility in the dimer. The highly abundant loop link (K622-E624) within the β3- αC loop of the monomer disappeared completely in the dimer and was replaced by a high-confidence intermolecular crosslink between K622 in the β3-αC loop and E668 in the αD helix. This observation is compatible with a face-to-face arrangement of the PKD kinase domain in the dimer, which has previously been observed experimentally for Chk2, the most closely related kinase to PKD for which a dimeric structure has been determined ^21^. Chk2 was therefore chosen as a template for comparative modeling of the PKD1 kinase domain dimer with experimental constraints derived from cross-linking mass spectrometry (XL-MS) using the Rosetta Comparative Modeling tool with C2 symmetry constraints ^22^. The resulting model revealed that the dimer interface additionally relies on a highly conserved acidic surface on the αG-helix consisting of E787, D788 and D791 (Supplementary Figure S3C) that forms an intricate network of salt bridges and hydrogen bonds with R741 and S742 in the activation loop as well as N757 in the αE/F-αF loop (Figure 1G). This is consistent with the stabilization of the activation loop indicated by XL-MS. To test this model, we introduced a single, charge-reversing mutation into the αG-helix (E787K), which completely abrogated dimerization of the kinase domain in solution (Figure 1C, H). Activation loop phosphorylation on S742 or the mutagenesis of S738 and S742 to glutamate (PKD^KD^ 2E) also prevents dimerization (Figure 1C, H) ^14^, presumably due to the introduction of negative charge in the vicinity of the acidic αG helix, which would lead to mutual repulsion. Single glutamate substitutions in either S738E or S742E are, however, not sufficient to dissociate the dimer (Figure 1C).

In summary, the kinase domain of PKD forms a stable face-to-face dimer in solution mediated by a network of highly conserved electrostatic interactions. Intriguingly, however, the hydroxyl group of S742 is sequestered 15.8 Å away from the gamma phosphate of ATP assuming a *trans*-reaction, and 13.1 Å away from the gamma phosphate of ATP assuming a *cis*-reaction, which makes this dimer incompatible with activation loop autophosphorylation as well as substrate binding. This observation prompted us to question what kinase domain dimerization is needed for in PKD1.

### Dimerization of the kinase domain is autoinhibitory

To quantitatively assess the auto-catalytic activity of the PKD kinase domain we performed a series of autophosphorylation assays. In order to capture the influence of dimerization, different concentrations of PKD1^KD^ were chosen in the range of 50 nM to 10 µM so as to cover the dynamic range of the monomer-dimer equilibrium (Figure 1C, black circles). Paradoxically for a reaction that was believed to occur in *trans*, autophosphorylation was more efficient with decreasing PKD concentration, observed reproducibly for PKD1^KD^ and PKD3^KD^ (Figure 2A). We therefore concluded that the kinase domains must dimerize in a manner that inhibits their ability to autophosphorylate. The affinity of homodimerization of the kinase domain on the basis of *trans*-autoinhibition was estimated to be 0.5-1.4 μM. To assess whether dimerization also affects substrate phosphorylation, we fused the homodimeric ULD of PKD1 (Figure 1C) to its kinase domain with a poly-glycine linker containing either five or ten glycine residues to force dimerization at a concentration (50 nM) at which the kinase domain alone is monomeric. Monitoring phosphate incorporation over time using a substrate peptide (Syntide2), we observed a five-fold decrease in the initial catalytic rate in both ULD-PKD1^KD^ chimeras, consistent with the formation of an inhibitory dimer (Figure 2B). Phosphorylation-induced dissociation of the kinase domains of PKD1 therefore has the potential to activate PKD simply by permitting substrate engagement.

**Figure 2.**
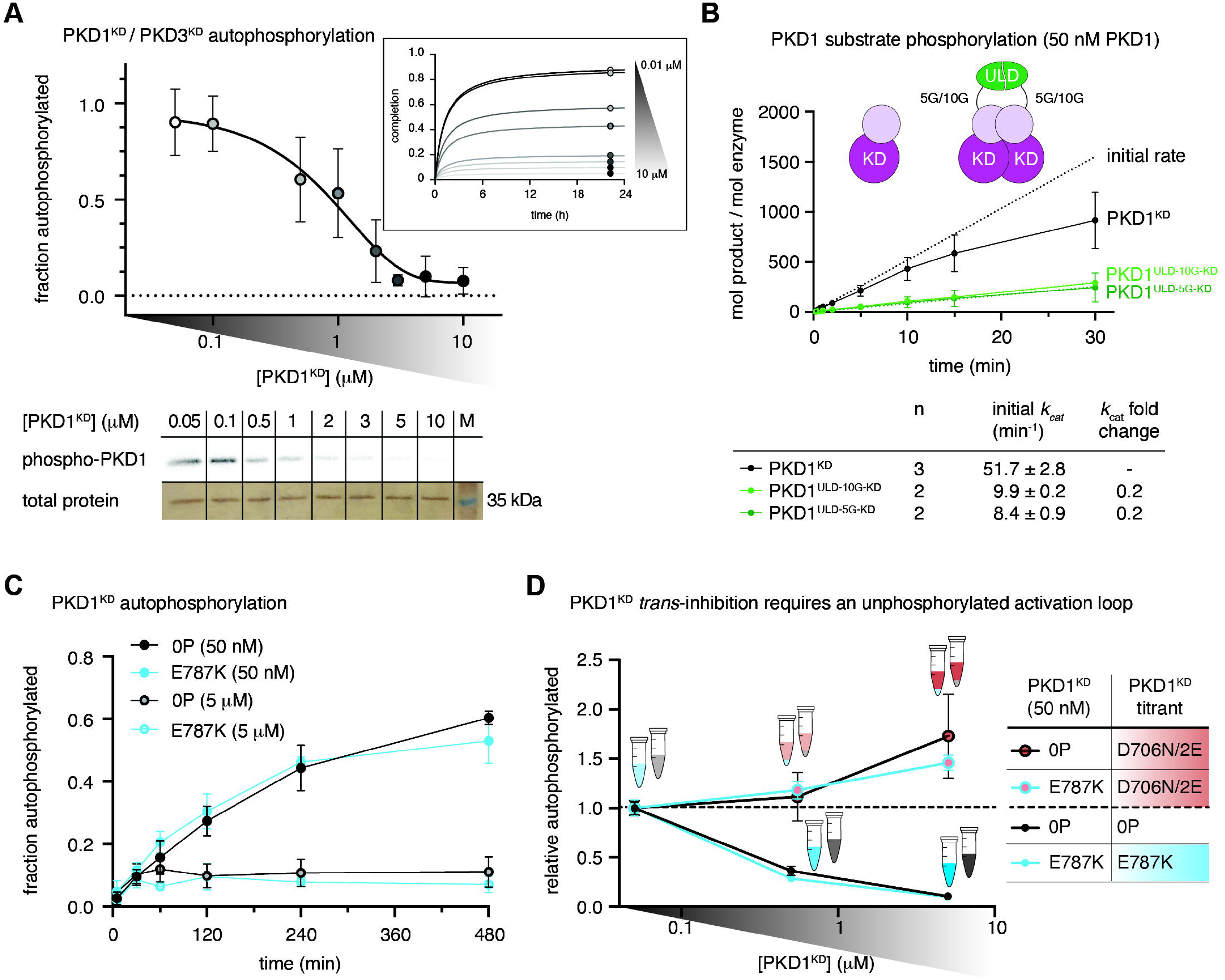
Dimerization of the kinase domain is autoinhibitory. A. Radiometric autophosphorylation assay of PKD1^KD^ and PKD3^KD^. Error bars are the standard deviation of 2-4 biologically independent experiments. Inset: data points represent reaction plateau at corresponding PKD1^KD^ concentrations. Bottom: representative autoradiograph of PKD1^KD^ samples compared to loading control (silver stain). B. Substrate phosphorylation kinetics of PKD1^KD^ compared to ULD-mediated PKD1^KD^ dimers with either (Gly)_5_ or (Gly)_10_ linkers between the ULD and kinase domains. Assay performed at concentrations corresponding to monomeric PKD1^KD^. Error bars are the standard deviation of *n* biologically independent experiments. Table indicates the number (n) of replicates and the values for *k*_cat_ derived from a linear regression in the linear range C. Autophosphorylation kinetics of wild-type PKD1^KD^ compared to monomeric PKD1^KD^ E787K at low (50 nM) and high (5 μM) concentrations. Error bars are the standard deviation of 3 biologically independent experiments. D. Autophosphorylation of wild-type PKD1^KD^ and PKD1^KD^ E787K in the presence or absence of catalytically dead, dimerization-incompetent PKD1^KD^ D706N/2E. Error bars are the standard deviation of 3 biologically independent experiments.

To further investigate the impact of dimerization on PKD activity, we measured the autophosphorylation and substrate phosphorylation kinetics of the previously identified PKD1^KD^ E787K interface mutant. The protein was monomeric by SEC-MALS (Figure 1H) and had no defect in substrate phosphorylation (Supplementary Figure S4A). Autophosphorylation at 50 nM was unaffected by the disruption of the interface (Figure 2C), which demonstrates that the dimer we observe in solution does not drive autophosphorylation. Surprisingly, however, increasing concentration of monomeric PKD1^KD^ E787K to 5 μM still resulted in inhibition of autophosphorylation that was indistinguishable from wild-type PKD1^KD^ (Figure 2C-D). This observation strengthens the notion that activation loop autophosphorylation does not occur in *trans*.

To further test whether molecular crowding could inhibit autophosphorylation by impeding productive collisions between PKD1^KD^ molecules, we made an inert kinase domain construct by combining a mutation of the catalytic aspartate (D706N) with S738E and S742E substitutions in the activation loop (PKD1^KD^ D706N/2E). This protein is dimerization deficient (Figure 1B), catalytically inactive (Supplementary Figure S4B), and cannot be phosphorylated. It is therefore an ideal mimic of PKD1^KD^ in both size and surface properties, while being unable to participate in the reaction. In contrast to both wild-type PKD1^KD^ and PKD1^KD^ E787K, increasing concentrations of PKD1^KD^ D706N/2E did not inhibit autophosphorylation of either PKD1^KD^ or PKD1^KD^ E878K (Figure 2D). These observations indicate that molecular crowding does not influence autophosphorylation, but rather that an unphosphorylated activation loop is required for *trans*-autoinhibition.

### PKD1 activation loop phosphorylation occurs in *cis*

Activation loop autophosphorylation in *cis* is highly controversial and little data exists to support its possibility. Nevertheless, our observations prompted us to ask whether indeed it could be the mechanism by which PKD activates itself. A *cis* reaction should be both concentration-independent and limited by the intrinsic enzymatic rate of phospho-transfer, which should result in linear kinetics until the reaction reaches stoichiometric completion. To test whether this is the case for PKD1, we analyzed the kinetics of autophosphorylation over a range of PKD1^KD^ concentrations at which the kinase domain is not *trans-*autoinhibited (10-50 nM). The kinetics of phosphate incorporation were linear at each concentration before reaching a plateau (Figure 3A) and, when normalized by the amount of PKD1^KD^ in the reaction, exhibited identical rates (Figure 3A). Since these findings strongly implied that the autophosphorylation reaction occurs in *cis*, we next asked whether or not wild-type PKD1^KD^ could phosphorylate a kinase-inactive PKD1^KD^ (D706N). Incubation of 10 nM wild-type PKD1^KD^ with an excess of 40 nM PKD1^KD^ D706N resulted in the incorporation of phosphate to a level that corresponds to 10 nM wild-type PKD1^KD^, and not to 50 nM total PKD1 (Figure 3B). To confirm that the phosphate was incorporated exclusively into wild-type PKD1^KD^ and not kinase-inactive PKD1^KD^, we analyzed the reaction by mass spectrometry. To distinguish between the two constructs, we incubated a shorter wild-type PKD1^KD^ truncated in its C-terminus by 18 amino acids (mass = 2078 Da) with PKD1^KD^ D706N at a ratio of 1:4 and a total concentration of 50 nM. Mass spectrometry of the resulting mixture revealed the incorporation of up to four phosphates (of which S742 is the primary modification^14^) exclusively into wild-type PKD1^KD^, but not kinase-inactive PKD1^KD^ (Figure 3C). These results unambiguously establish that activation loop autophosphorylation of PKD1 occurs in *cis* and not in *trans*.

**Figure 3.**
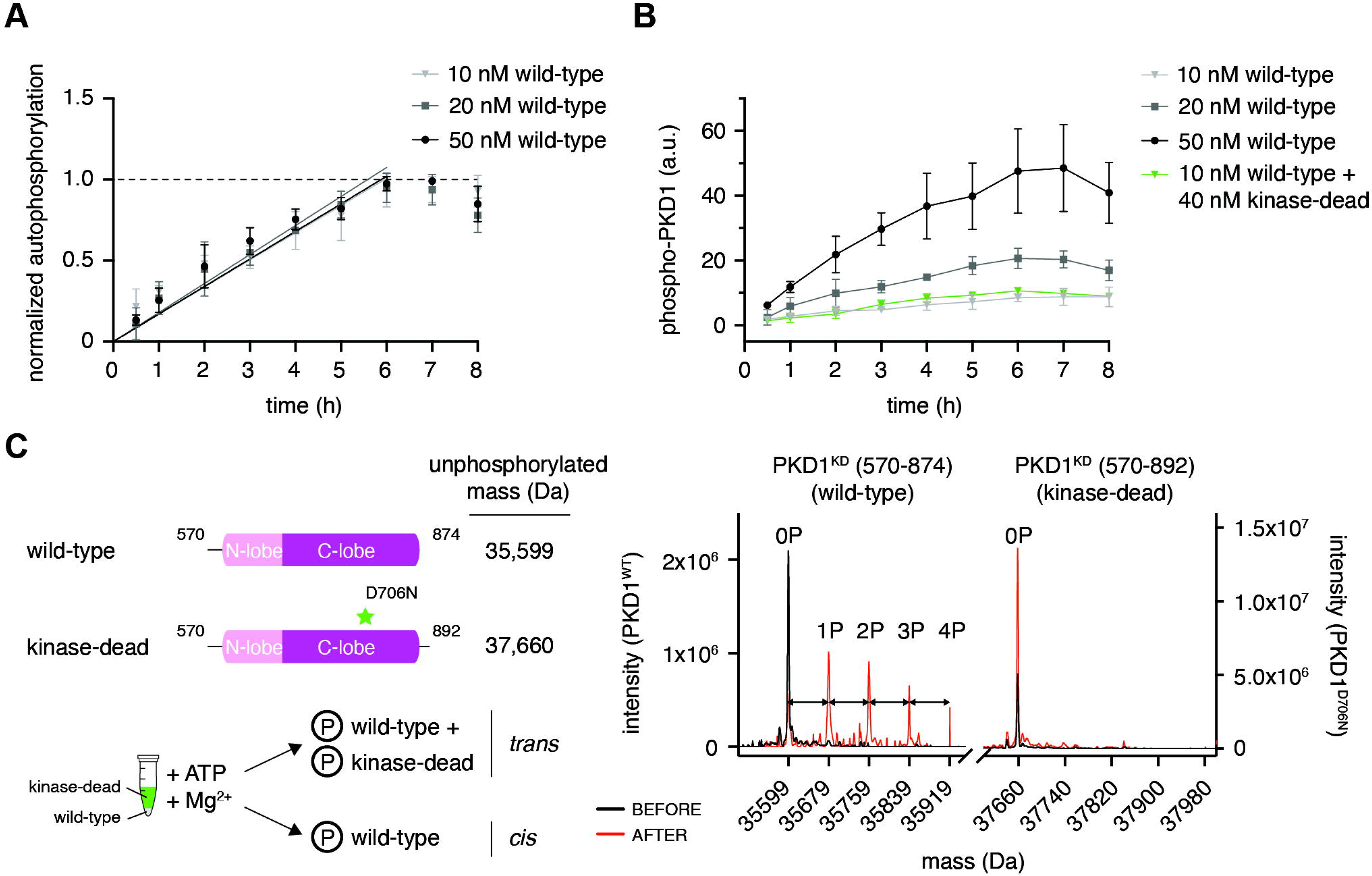
PKD1 activation loop autophosphorylation occurs in *cis*. A. Activation loop autophosphorylation kinetics of PKD1^KD^ at low concentration. Error bars are the standard deviation of 3 biologically independent experiments. B. Activation loop autophosphorylation of kinase-dead PKD1^KD^ D706N in the presence and absence of excess wild-type PKD1^KD^. Error bars are the standard deviation of 3 biologically independent experiments. C. Mass spectrometry analysis of the autophosphorylation reaction containing 10 nM wild-type PKD1^KD^ and 40 nM kinase-dead PKD1^KD^ D706N shown in panel B. Phospho-species of wild-type PKD1^KD^ are separated by 80 Da (double-headed arrows).

### Activation loop phosphorylation increases PKD catalytic activity

Activation loop phosphorylation has long been identified as a crucial step in the activation of PKD ^23^, but the effect of this regulatory modification at the molecular level was only partially understood ^24^. We have shown in this (Figure 1B) and previous work ^14^ that activation loop phosphorylation on S742 prevents kinase domain dimerization, which, in the context of an inhibitory kinase domain dimer, may be sufficient to activate PKD simply by relieving autoinhibition. Whether phosphorylation also increases the intrinsic catalytic activity of PKD is, however, unknown. To dissect the structural impact of activation loop phosphorylation used hydrogen-deuterium exchange mass spectrometry (HDX-MS) to compare the dynamics of PKD1^KD^ 0P to PKD1^KD^ 2E, both in the presence of ATP and Mg^2+^. In PKD1^KD^ 2E compared to PKD1^KD^ 0P we observed a strong and very specific decrease in the rates of HD exchange in the activation loop and the catalytic loop as well as the αC helix, β1 and β3 strands (Figure 4A-B, Supplementary Table S2, Supplementary Figure S5A). Together, these elements constitute the kinase active site, and their stabilization is consistent with the stereotypical role of activation loop phosphorylation. Correspondingly, the substrate phosphorylation rates of both stoichiometrically S742-phosphorylated PKD1^KD^ (1P) and PKD1^KD^ 2E were 3-fold higher compared to unphosphorylated PKD1^KD^ (0P) (Figure 4C).

**Figure 4.**
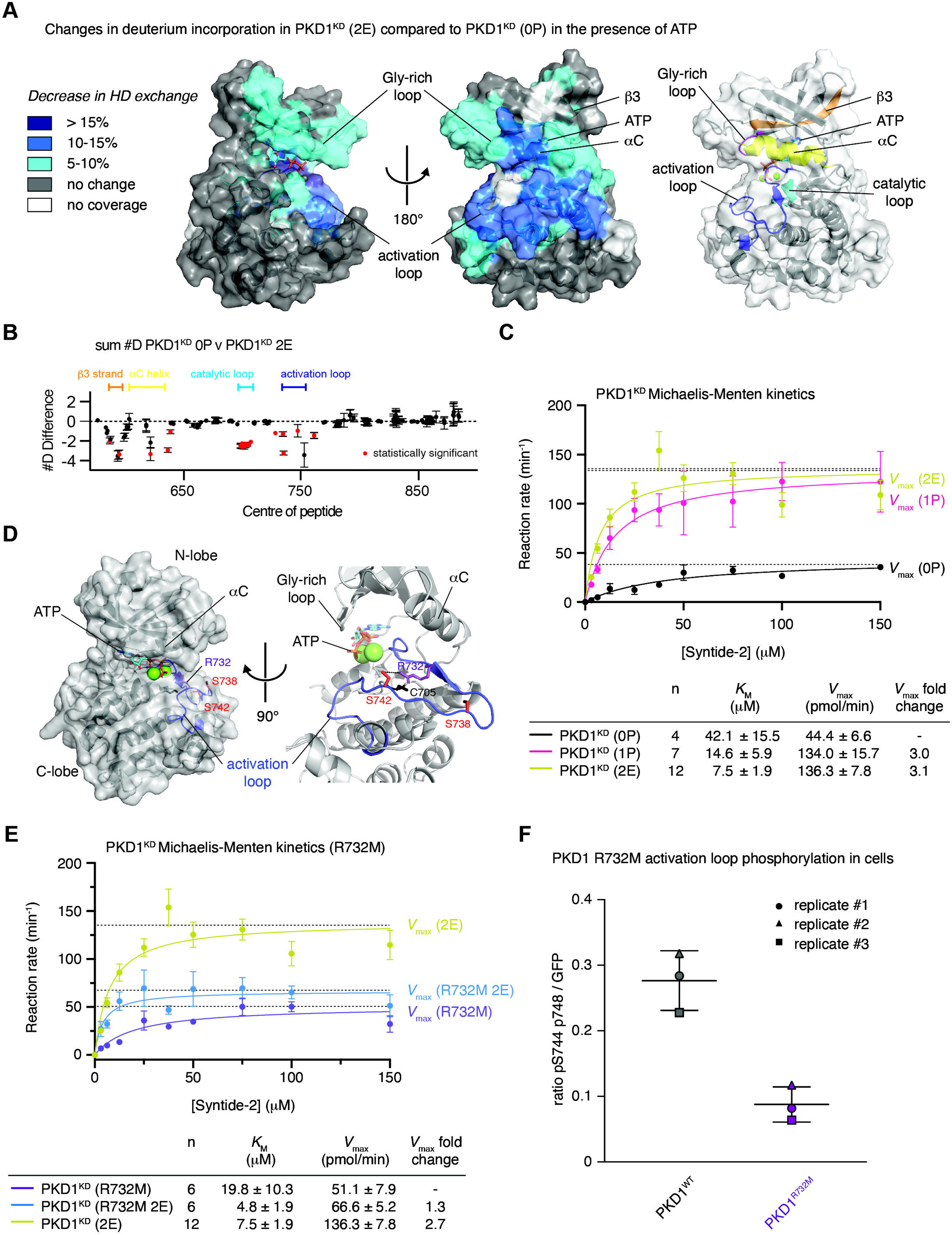
Activation loop autophosphorylation increases PKD1 catalytic activity. A. Significant differences in deuterium incorporation in PKD1^KD^ (2E) compared to unphosphorylated PKD1^KD^ (0P) in the presence of ATP. Changes mapped onto the kinase domain of PKD1. Differences in exchange in a peptide were considered significant if they met all three of the following criteria: ≥5% change in exchange and a ≥0.5 Da difference in exchange with a 2-tailed T-test value of less than 0.01 at any timepoint. Right: color-coded cartoon of key structural elements that exhibit significant changes. Glycine-rich loop (magenta), strand β3 (orange), helix αC (yellow), catalytic loop (cyan), activation loop (blue). B. The number of deuteron difference for all peptides analyzed over the entire deuterium exchange time course is shown, with each point representing an individual peptide (full exchange information for every peptide available in the source data). Statistically significant changes are indicated in red. C. Michaelis-Menten kinetic analysis of PKD1^KD^ substrate phosphorylation. Unphosphorylated PKD1^KD^ (0P) (black), S742-phosphorylated PKD1^KD^ (1P) (magenta), PKD1^KD^ (2E) (yellow-green). Error bars are the standard deviation of *n* biologically independent experiments. Table indicates the number (*n*) of independent biological replicates and the values for *K*_M_ and *V*_max_ derived from fitting the data with the Michaelis-Menten equation. D. Rosetta homology model of the PKD1 kinase domain, indicating the arrangement of side chains surrounding S742 in the activation loop. Activation loop (blue), phospho-acceptor residues (red). E. Michaelis-Menten enzyme kinetics of PKD1^KD^ R732M (purple) compared to PKD1^KD^ (2E) (yellow-green) and PKD1^KD^ R732M 2E (marine blue). Error bars are the standard deviation of *n* biologically independent experiments. Table indicates the number (*n*) of independent biological replicates and the values for *K*_M_ and *V*_max_ derived from fitting the data with the Michaelis-Menten equation. F. PKD1 activation loop phosphorylation in HEK293T cells. Wild-type PKD1 (black), PKD1 R732M (purple).

Typically, kinases that depend on activation loop phosphorylation to acquire full activity rely on a conserved arginine in the so-called HRD motif to make electrostatic interactions with the respective phosphate group and thereby stabilize the activation loop in in its active conformation. PKD, however, encodes a cysteine at this position (Supplementary Figure S5B), which excludes it from the classical group of RD kinases. Whilst it has been postulated that only RD kinases can be regulated by activation loop phosphorylation ^6, 7^, *in silico* modeling (Rosetta) of the PKD1^KD^ monomer revealed that R732 projects its guanidinium group into precisely the same three-dimensional position as the arginine in classical RD kinases (Figure 4D, Supplementary Figure S5C). Mutation of R732 to methionine in PKD1^KD^ abrogated the increase in catalytic activity gained by phosphorylation or phosphomimetic substitutions (Figure 4E), while the substrate phosphorylation kinetics of unphosphorylated PKD1^KD^ R732M were not significantly different to unphosphorylated, wild-type PKD1^KD^ (0P) (Figure 4C). Taken together, these data indicate that R732 in PKD1 plays an analogous role to the arginine of the classical HRD motif of eukaryotic protein kinases. When overexpressed in HEK293 cells, PKD1^R732M^ exhibited dramatically reduced activation loop phosphorylation, presumably due to enhanced dephosphorylation as a consequence of the loss of coordination of the phosphate (Figure 4F, Supplementary Figure S6).

In summary, activation loop autophosphorylation activates PKD in two distinct ways: first, by preventing autoinhibitory dimerization of its kinase domain and, second, by increasing its intrinsic catalytic activity.

### PKD auto- and substrate phosphorylation are mechanistically distinct

Substrates of PKD share a conserved consensus sequence (Syntide-2) that is not exhibited by either the phosphorylation motifs (S738 and S742) in the activation loop or an additional autophosphorylation motif in the C-terminus of PKD1 and PKD2 (Figure 5A). Kinases of the CAMK and AGC families specifically recognize an arginine in the P-3 position by virtue of a salt bridge to a conserved aspartate or glutamate residue (D665 in PKD1) in the β5-αD loop (Figure 5A) ^25^. Mutation of D665 to asparagine (D665N) in PKD1 was previously reported to result in a loss of substrate specificity and consequent oncogenic rewiring, but not a loss of activity ^26^. Purified, recombinant PKD1^KD^ D665N, however, was completely inactive against the PKD consensus substrate peptide Syntide-2 *in vitro* (Figure 5B), and the activity of full-length PKD1^D665N^ against a Golgi-localized PKD activity reporter GPKDrep ^27^ was significantly reduced compared to wild-type PKD1 and similar to kinase-dead PKD1^K612W^ in cells (Figure 5C). In vitro autophosphorylation of PKD1^KD^ D665N, on the other hand, was indistinguishable from wild-type PKD1^KD^ (Figure 5D). We next investigated phosphorylation of PKD1^D655N^ in cells under basal conditions and after stimulation by nocodazole, which is known to strongly increase PKD activity ^28^. Interestingly, S738 and S742 in the activation loop as well as pS910 in the C-terminus of PKD1 were hyperphosphorylated in PKD1^D665N^ in cells when compared to wild-type PKD1 and PKD1^K612W^ under basal conditions and with nocodazole stimulation. Notably, S910 phosphorylation, which was absent in PKD1^K612W^, increased further upon stimulation for wild-type PKD1 but not for PKD1^D665N^ (Figure 5E-F, Supplementary Figure S7A-B).

**Figure 5.**
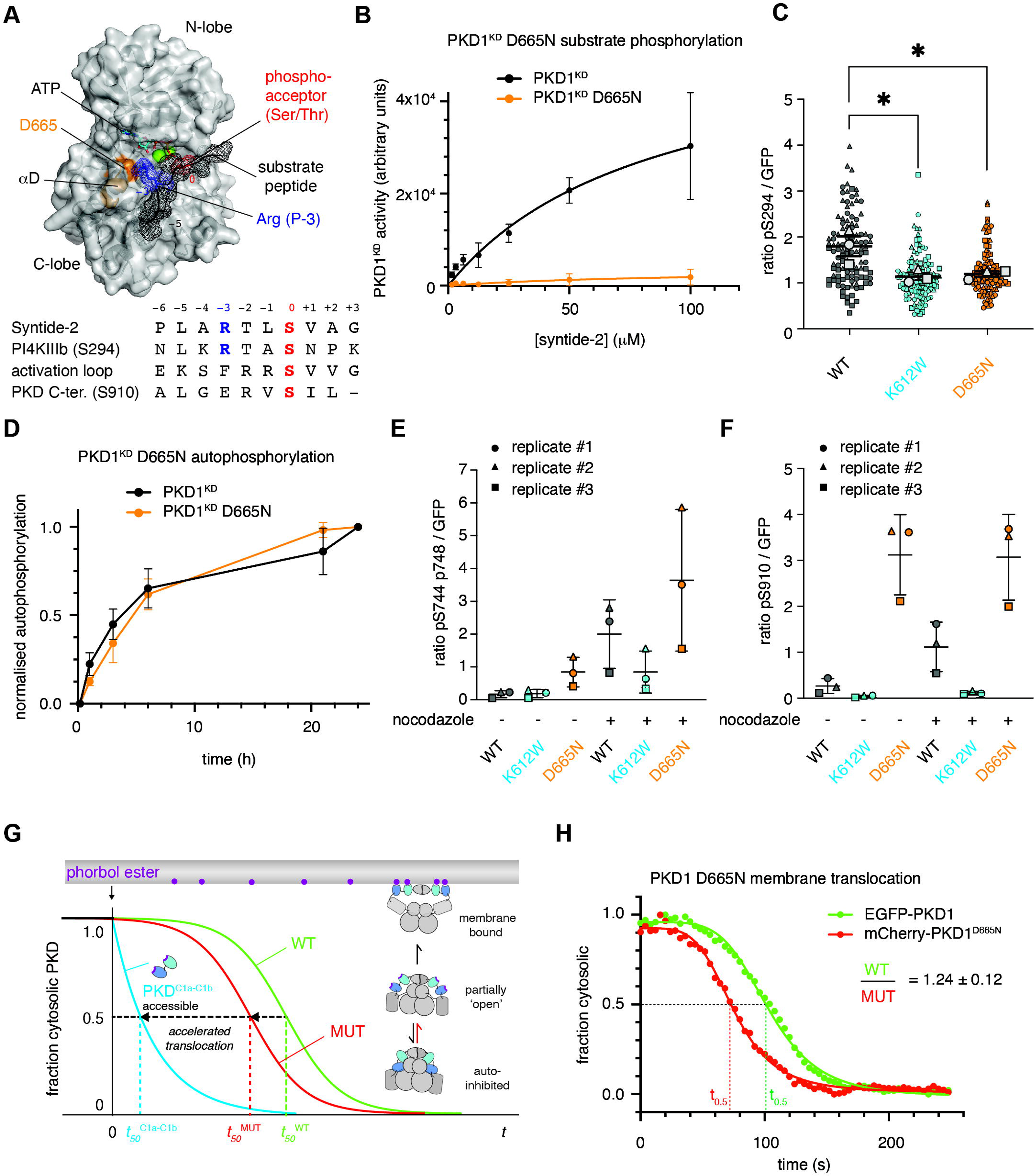
PKD1 auto- and substrate phosphorylation are mechanistically distinct. A. Homology model of the PKD1^KD^-substrate interaction. Substrate peptide (black mesh), phospho-acceptor of substrate (red), Arg at P-3 position (blue), residue D665 in helix αD (orange). Below: alignment of consensus sequence (Syntide-2), the PKD substrate PI4KIIIβ (S294), the PKD activation loop sequence (S742), and the C-terminal autophosphorylation motif (S910). B. PKD1^KD^ (black) substrate phosphorylation kinetics compared to PKD1^KD^ D665N (orange). Error bars are the standard deviation of 2 biologically independent experiments. C. PKD1-mediated GPKDrep phosphorylation in HeLa cells determined by ratiometric imaging. Wild-type PKD1 (black), PKD1 K612W (blue), PKD1 D655N (orange). Data are represented in symbol-coded bee swarm SuperPlots, with each cell-level data point revealing which experiment it came from. Lines show mean and SEM of the replicate means. Statistical test: ordinary one-way ANOVA with Sidak’s multiple comparison test, *p < 0.05. D. Autophosphorylation kinetics of wild-type PKD1^KD^ (black) compared to PKD1KD D665N (orange). Error bars are the standard deviation of 3 biologically independent experiments. E. PKD1 activation loop phosphorylation in HEK293T cells determined by western blot analysis. Wild-type PKD1 (black), PKD1 K612W (blue), PKD1 D655N (orange). Lines show mean and SEM of the replicate means. F. PKD1 S910 phosphorylation in HEK293T cells determined by western blot analysis. Wild-type PKD1 (black), PKD1 K612W (blue), PKD1 D655N (orange). Lines show mean and SEM of the replicate means. G. Cartoon schematic of PKD membrane translocation assay. The tandem DAG-binding domains (blue) translocate rapidly to the plasma membrane upon phorbol ester treatment. Mutations that destabilize the intramolecular assembly of PKD accelerate its membrane translocation kinetics (red) compared to wild-type PKD (green). H. Membrane translocation kinetics of wild-type EGFP-PKD1 (green) compared to mCherry-PKD1^D665N^ (red) in Cos7 cells. Ratio of times for half-maximal translocation derived from analysis of 3 cells.

Since activation loop autophosphorylation depends on dissociation of the inhibitory kinase domain dimer, we speculated that D665N may weaken the intramolecular assembly in the context of full-length PKD1, thereby leading to activation loop hyperphosphorylation. To test whether this is the case, we employed a ratiometric membrane translocation assay (Figure 5G) in which we measure the rates of translocation of wild-type and mutant PKD variants to the plasma membrane upon treatment of the cells with phorbol ester, a natural DAG mimetic. Since the binding to phorbol esters is essentially irreversible, the rate of translocation is correlated with the accessibility of the phorbol ester-binding C1 domains. By co-transfecting spectrally separable fluorescent fusion proteins, wild-type and mutant PKD variants can be compared within the same cell ^29^. In the autoinhibited conformation of PKD, the membrane binding surfaces of the C1 domains are buried in intramolecular interactions ^14^. PKD1^D665N^ translocated to the plasma membrane significantly faster than wild-type PKD1 (Figure 5H), indicating that the mutation destabilizes the autoinhibited conformation.

In summary, these observations unambiguously confirm that activation loop *cis*- autophosphorylation and *trans*-phosphorylation of downstream substrates by PKD are mechanistically distinct, and that D665 plays an essential role in both substrate phosphorylation and PKD1 autoinhibition.

### Substrate binding-deficient and constitutively dimeric PKD1 block secretion

PKD controls the fission of basolateral membrane-targeted cargo vesicles from the TGN ^9^. To investigate the impact of the substrate phosphorylation-deficient D665N mutant on protein secretion we monitored the distribution of pancreatic adenocarcinoma upregulated factor (PAUF), a known PKD-dependent cargo. PAUF secretion is arrested in the TGN by overexpression of kinase-dead PKD^K612W^ ^9^. Consistent with the loss of substrate phosphorylation, but not autophosphorylation, we observed the same Golgi accumulation of PAUF in cells transfected with PKD1^D665N^ (Figure 6A), while PAUF secretion was completely abrogated (Figure 6B). This further strengthens the essentiality of PKD in secretion, wherein it controls the fission of cargo vesicles from the TGN.

**Figure 6.**
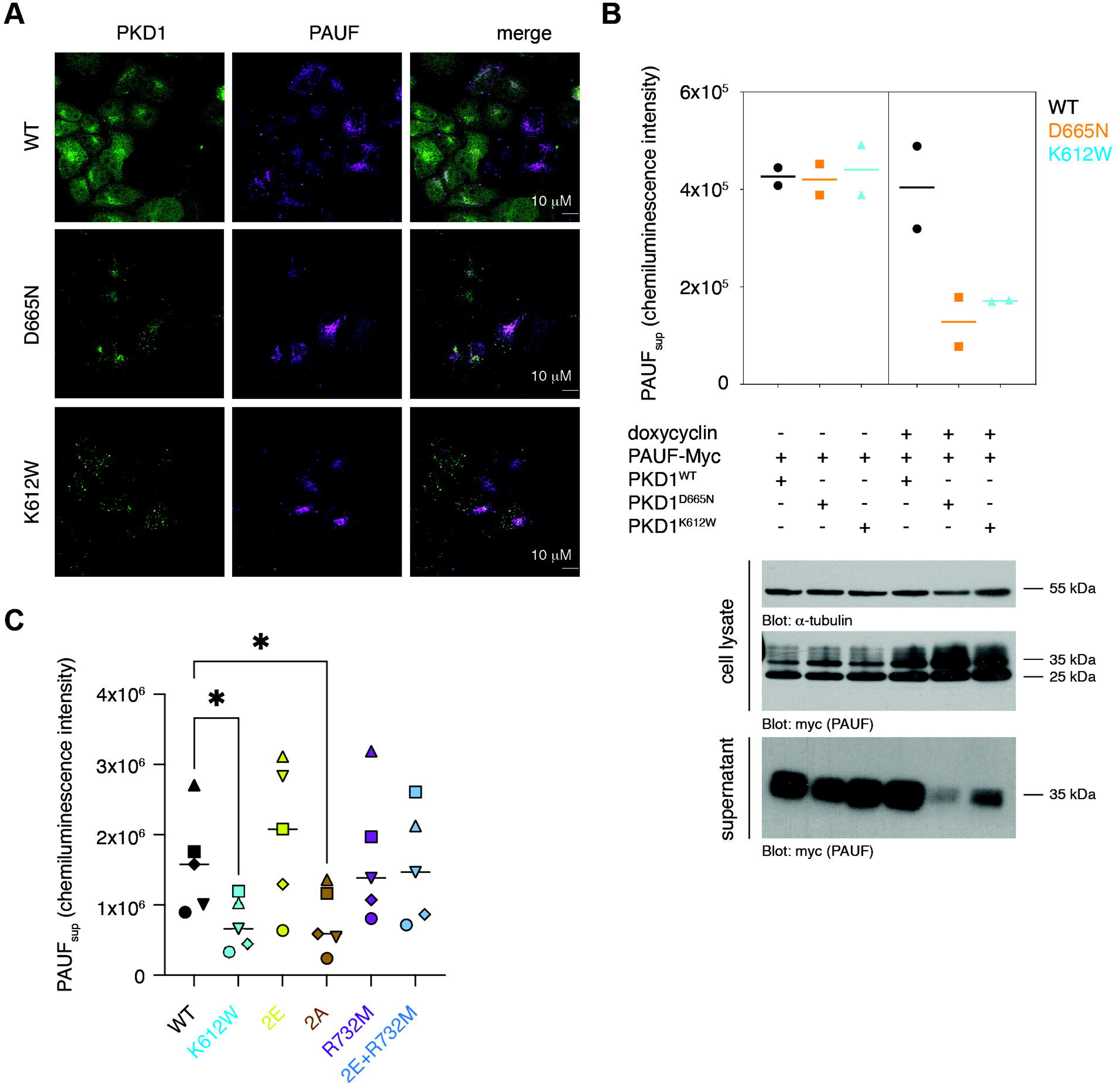
Substrate binding-deficient and constitutively dimeric PKD1 block secretion. A. Subcellular localization of PKD1-GFP variants and PAUF in HeLa cells. B. PAUF secretion in FlpIN-TRex-HeLa cells under conditions of stable doxycycline-inducible PKD1 expression and transient constitutive PAUF expression. Lines show mean of replicate data. C. PAUF secretion in HEK293T cells under conditions of transient constitutive PKD1 and PAUF expression. Lines show mean of replicate data. Statistical test: RM one-way ANOVA with Sidak’s multiple comparison test, *p < 0.05.

We previously demonstrated that activation loop phosphorylation prevents kinase domain dimerization, thereby relieving autoinhibition in addition to increasing catalytic activity (Figure 1C, 4C). Conversely, a mutant of the kinase domain in which both phosphorylation sites are mutated to alanine (PKD1^KD^ 2A), is dimeric in vitro (Figure 1C). This implies that PKD that is unable to be phosphorylated might exert a dominant negative effect on secretion in cells due to constitutive autoinhibition. Indeed, PKD1^2A^ abrogated PAUF secretion (Figure 6C, Supplementary Figure S8), despite the fact that unphosphorylated PKD1^KD^ (0P) has substantial basal activity *in vitro* (Figure 4C) and the protein is ectopically overexpressed. In contrast, expression of PKD1^R732M^ does not impair PAUF secretion (Figure 6C-D), presumably due to low, but not absent, activation loop phosphorylation in stimulated cells (Figure 4F) and reduced, but not abrogated, activity when combined with activation loop phosphomimetics in vitro (Figure 4E). As such, the reduced activity of PKD1^R732M^ is likely compensated for by overexpression. Together, these observations strengthen the physiological relevance of a face-to-face autoinhibitory dimer of the kinase domains in the regulation of PKD activity.

### Autoregulation of membrane binding depends on ATP, but not dimerization

Previous work has established that the DAG-binding C1 domains of PKD are sequestered in its autoinhibited, cytosolic conformation ^14^. The endogenous concentration of PKD has been estimated to be 3-32 nM ^14, 30^. Dimerization of PKD via both its ULD and kinase domains, which have affinities of homodimerization of 0.5-2 µM and 0.9 µM respectively, implies that PKD is constitutively dimeric in cells and autoinhibited in the absence of DAG. To test whether dimerization is relevant for the regulation of DAG binding, we once again examined membrane translocation kinetics as a proxy for the stability of the autoinhibited state. Consistent with PKD being dimeric in the cell, fusion of the parallel coiled-coil of the yeast transcription factor GCN4 to the N-terminus of PKD1 did not affect membrane translocation in response to phorbol ester stimulation (Figure 7A). Disruption of kinase domain dimerization either by introduction of phosphomimetics in the activation loop (2E) or mutation of the αG interface (E787K), resulted in membrane translocation kinetics that were indistinguishable from PKD1^WT^ (Figure 7B-C). Finally, we combined a mutation in the ULD dimer interface (F104E in PKD1, F59E in CeDKF1) (Figure 1C) ^14^ with the kinase domain 2E mutant (PKD1^F104E+S738E/S742E^) to create a construct which is compromised in all known dimerization interfaces. This construct, which is presumably monomeric, also did not show altered membrane translocation kinetics (Figure 7D). We therefore conclude that the autoregulation of DAG binding does not intrinsically depend on PKD dimerization.

**Figure 7.**
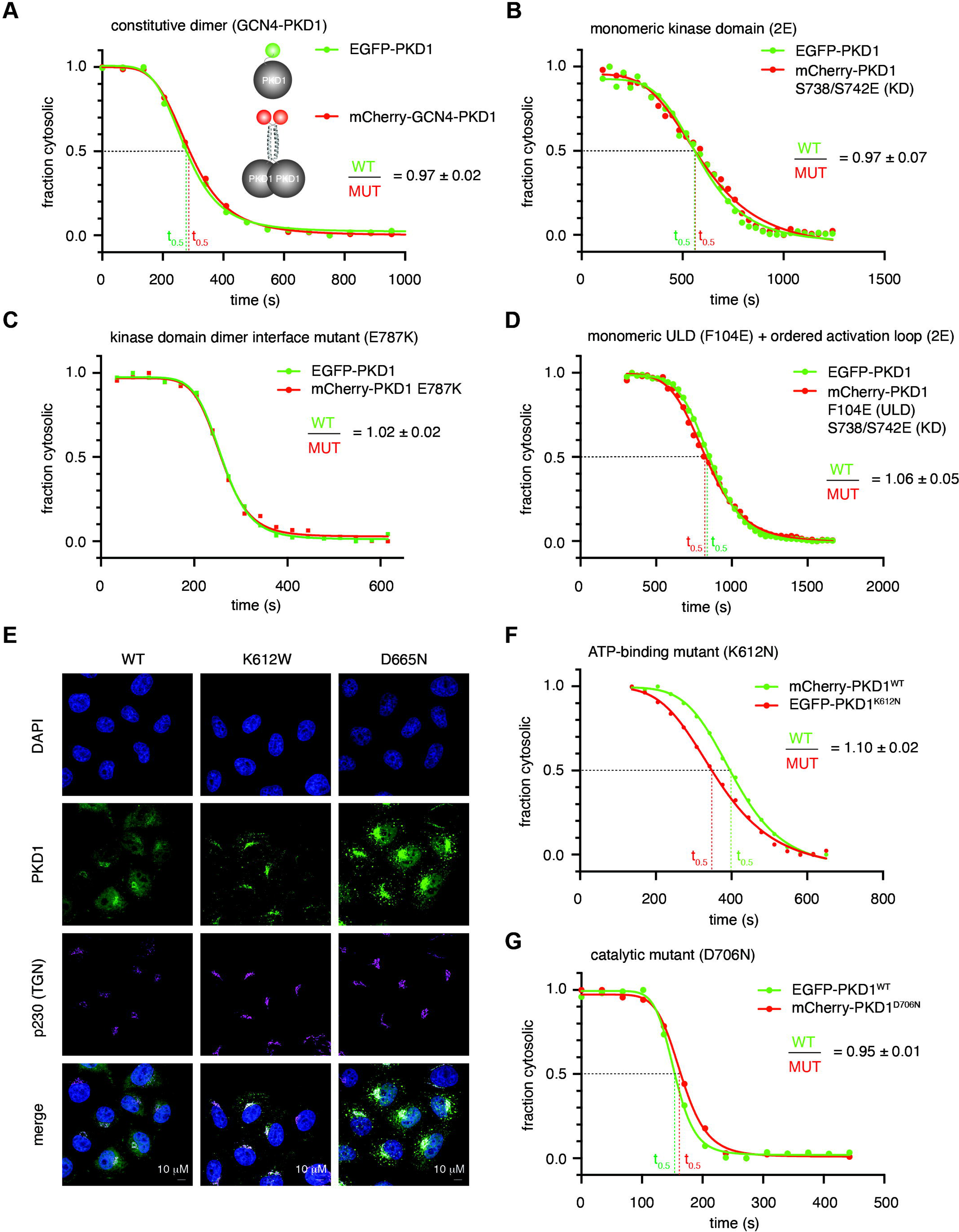
Autoregulation of membrane binding depends on ATP, but not dimerization. A. Membrane translocation kinetics of GCN4-PKD1 (mCherry, red) compared to wild-type PKD1 (EGFP, green) in Cos7 cells. Ratio of times for half-maximal translocation derived from analysis of 5 cells. B. Membrane translocation kinetics of PKD1^2E^ (mCherry, red) compared to wild-type PKD1 (EGFP, green) in Cos7 cells. Ratio of times for half-maximal translocation derived from analysis of 4 cells. C. Membrane translocation kinetics of PKD1^E787K^ (mCherry, red) compared to wild-type PKD1 (EGFP, green) in Cos7 cells. Ratio of times for half-maximal translocation derived from analysis of 4 cells. D. Membrane translocation kinetics of PKD1^F104E+2E^ (mCherry, red) compared to wild-type PKD1 (EGFP, green) in Cos7 cells. Ratio of times for half-maximal translocation derived from analysis of 4 cells. E. Subcellular localization of wild-type PKD1-GFP, PKD1^K612W^-GFP (ATP binding mutant) and PKD1^D665N^-GFP (substrate binding mutant) in stable Flp-In TRex HeLa cells. F. Membrane translocation kinetics of PKD1^K612N^ (EGFP, red) compared to wild-type PKD1 (mCherry, green) in Cos7 cells. Ratio of times for half-maximal translocation derived from analysis of 3 cells. G. Membrane translocation kinetics of PKD1^D706N^ (mCherry, red) compared to wild-type PKD1 (EGFP, green) in Cos7 cells. Ratio of times for half-maximal translocation derived from analysis of 5 cells.

Previous studies have shown an enrichment of kinase-dead PKD at the TGN when compared to wild-type PKD ^9^, which we could recapitulate. PKD1^D665N^ also exhibited a similar enrichment at the TGN, consistent with its sensitivity to phorbol esters (Figure 7E, Figure 5H). To assess the mobility of PKD1 in real time, we performed photobleaching experiments to measure the fluorescence recovery after photobleaching (FRAP) of individual TGNs from PKD1^WT^, PKD1^K612W^, and PKD1^D665N^ -GFP expressing Hela cells. To quantitatively measure PKD1 localization to the TGN, FRAP data were individually fitted with a one-phase exponential function to obtain values for the mobile fraction and the rate of fluorescence recovery (half-time of recovery). While wild-type PKD1 exhibited fast fluorescence recovery (10,52s (95% CI 8,644 to 12,57)), the half-time of PKD1^K612W^ (53.7s (95% CI 45,93 to 63,30)) and PKD1^D665N^ (39.34s (95% CI 36,09 to 42,91)) recovery was strongly increased demonstrating increased residency time for both mutants at the TGN. Moreover, the mobile fraction of PKD1^D665N^ was significantly reduced compared to wild-type PKD1, showing that PKD1^D665N^ associates tightly with the TGN membrane (Supplementary Figure S9A-C). Since K612W and D665N mutations impair ATP and substrate binding respectively, we asked whether ATP binding is important for the stability of the autoinhibited conformation of PKD and the sequestration of its membrane binding C1 domains. Mutation of K612 resulted in accelerated translocation kinetics (Figure 7F) comparable to D665N (Figure 5H). Intriguingly, however, mutation of the catalytic aspartate in the active site to asparagine (D706N) had no effect on membrane binding (Figure 7G). These results indicate that ATP binding, but not catalysis, is important for stabilization of the intramolecular, autoinhibited assembly.

In summary, the stable intramolecular assembly of the regulatory and kinase domains of PKD1 depends on ATP binding but is independent of catalysis and dimerization. Disruption of ATP binding leads to premature membrane translocation, presumably by destabilizing the autoinhibited conformation, and accumulation of PKD at the TGN. This is also consistent with inhibitory kinase domain dimerization since DAG-mediated *cis*-autophosphorylation of PKD restricts PKD activity to the TGN.

## Discussion

The majority of eukaryotic protein kinases that auto-activate by activation loop phosphorylation are believed to do so in *trans*. This necessitates their transient dimerization. In this study, we demonstrate that kinases can be regulated by the inverse mechanism: dissociation of an inhibitory dimer followed by autophosphorylation in *cis* (Figure 8). Whilst our study focuses specifically on the regulation of PKD, *cis*- autophosphorylation of its activation loop is likely to be a much more common mode of kinase regulation.

**Figure 8.**
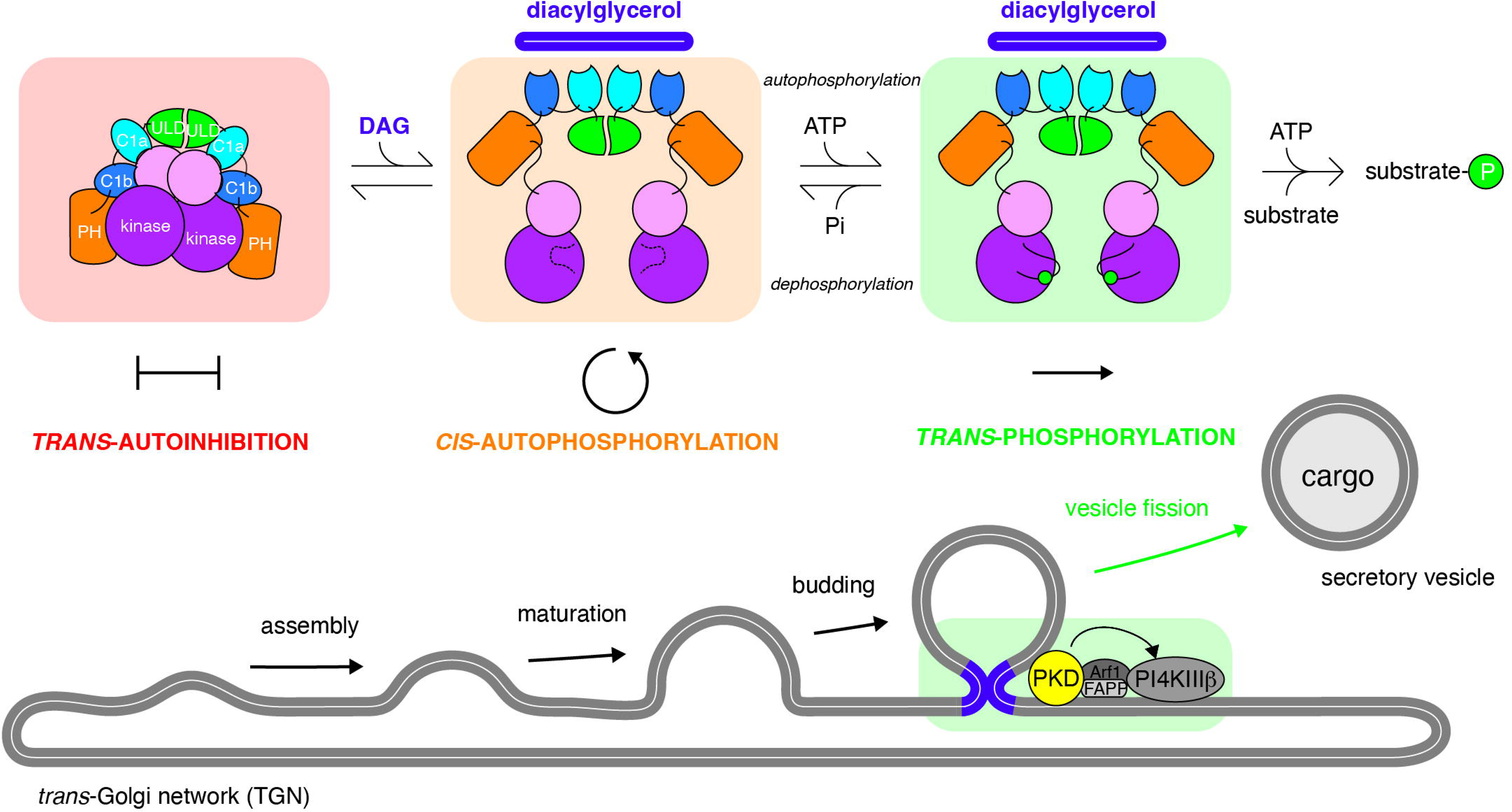
PKD autoinhibition in *trans* regulates activation loop autophosphorylation in *cis*. Model for the control of PKD by DAG binding and subsequent triggering of cargo vesicle fission from the TGN. PKD adopts a constitutively dimeric conformation in the cytosol of cells in which its kinase domains are maintained in a trans-autoinhibitory face-to-face dimer. Binding to DAG at the TGN, presumably in the vicinity of the bud neck of cargo vesicles, results in conformational changes that relieve the inhibitory dimerization of its kinase domains, leading to activation loop *cis*-autophosphorylation. Activation loop phosphorylation both promotes *trans*-phosphorylation of substrates and prevents reassembly of the kinase domains.

Our findings lead to a revised model of PKD regulation (Figure 8). In the absence of activating signals, PKD is maintained in a dimeric, autoinhibited conformation mediated by homodimerization of its ULD and kinase domains. The inactive conformation is characterized by a face-to-face arrangement of its kinase domains that inhibits both activation loop autophosphorylation and substrate phosphorylation. The inactive conformation of PKD sequesters its regulatory C1 domains in intramolecular interactions that inhibit binding to DAG in the membranes of the TGN. Upon DAG enrichment in the TGN, the C1 domains disengage from the intramolecular assembly, triggering conformational changes that are favorable for activation loop autophosphorylation in *cis*. Once phosphorylated, the kinase domain of PKD acquires full catalytic activity and is unable to dimerize, thereby protecting it from *trans*-autoinhibition. PKD is thereby fully primed to initiate downstream signaling, resulting in cargo secretion from the TGN.

Autoinhibition is common to many protein kinases and serves to inactivate them in the absence of a stimulus. The αG helix is at the center of a surface epitope commonly employed in the regulation of kinase activity. This includes the association of regulatory domains ^31, 32^, the binding of substrates ^33^, the recruitment of inactivating phosphatases^34^, or the association of kinase domains to catalyze *trans*-autophosphorylation ^35–37^. Face-to-face dimers of the related death associated protein kinases (DAPK) have previously been observed ^38, 39^, yet these kinases do not autophosphorylate their activation loop. Their homodimerization is stably maintained by a constitutive leucine zipper coiled-coil domain. The face-to-face dimerization of their kinase domains serves to maintain them in an autoinhibitory conformation ^38, 39^, which in DAPK2 is relieved by the binding of calmodulin to the C-terminal autoregulatory domain ^38^. Truncation of the coiled-coil domain of Zipper-interacting protein kinase (ZIPK) results in its activation ^40^, suggesting that it helps mediate autoinhibition. Monomeric DAPKs maintain their activation loop in the active conformation not via phosphorylation, but via a unique set of hydrophobic interactions between conserved phenylalanines in the activation loop and the C-lobe ^41^. In PKD, the dissociation of the kinase domains is required in order to accomplish activation loop *cis*-autophosphorylation.

PKD activation loop phosphorylation has long been thought to occur in *trans* on the basis that catalytically inactive PKD expressed in cells acquires S738/S742 activation loop phosphorylation, presumably via endogenous PKD ^13^. Our own recent discovery of the ULD dimerization domain in the *C. elegans* PKD homolog, Dkf1, led us to conclude that this mediates phosphorylation in *trans* ^14^. Activation loop autophosphorylation in *cis* is widely considered not to be possible on the basis that the activation loop runs in the opposite direction to a canonical substrate peptide, and that the inactive conformation of the activation loop has often been observed to fold back into the active site, disrupting the catalytic machinery. While some kinases have been reported to autophosphorylate in *cis* via a variety of mechanisms including the formation of translational and chaperone-assisted intermediates ^42^, rationalization of such mechanisms at the atomic level is still missing. However, structures of the insulin receptor kinase (IRK) and fetal liver tyrosine kinase (FLT3) provide compelling evidence that a *cis* arrangement of the activation loop in a phosphorylation-competent conformation is physically possible ^43, 44^. The activation loop tyrosine, which undergoes autophosphorylation in both of these kinases, superimposes perfectly with the phospho-acceptor tyrosine of a substrate peptide ^45^.

We have shown that the mechanisms of autophosphorylation and substrate phosphorylation in PKD are mechanistically distinct: indeed, we demonstrate that activation loop autophosphorylation occurs exclusively in *cis*. However, the kinetics of PKD *cis*-autophosphorylation are extremely slow (Figure 2C, Figure 3C), suggesting that PKD relies on additional activating inputs that are as yet unknown. Since activation loop autophosphorylation prevents autoinhibition in *trans*, the reaction must be prohibited in the absence of an activating stimulus. In the context of a *cis*-reaction that must be prevented unless coupled to DAG binding, the slow kinetics perhaps safeguard the kinase domain from spurious activation during transient dissociation events. The enhanced *cis*-autophosphorylation reaction rates elicited by GST-mediated dimerization demonstrate that the intrinsic reaction rate of the monomeric kinase domain can be upregulated. As such, identification of the molecular trigger that promotes activation loop phosphorylation post-dissociation of the kinase domains will be important.

The closest related kinase to PKD, Checkpoint-associated kinase 2 (Chk2), has been reported to autophosphorylate its activation loop in *trans*. Whereas the DAPKs are obligate homodimers by virtue of their C-terminal coiled-coil and PKD is likely an obligate ULD-mediated homodimer, Chk2 undergoes dimerization via a forkhead-associated (FHA) domain that depends on phosphorylation of Thr68 by the ataxia telangiectasia mutated (ATM) kinase ^46–49^. The current model of Chk2 regulation supposes that T68 phosphorylation by ATM in response to DNA-damage leads to cell cycle checkpoint activation via T68-mediated Chk2 dimerization and *trans*- autophophorylation. The structure of a construct of Chk2 containing both its FHA and kinase domains revealed that FHA-mediated dimerization of Chk2 drives face-to-face apposition of its kinase domains in an arrangement that was interpreted to represent the *trans*-autophosphorylation reaction ^21^. A similar structure has been obtained for the yeast homolog of Chk2, Rad53 ^50^. GST-mediated dimerization of the Chk2 kinase domain has previously been shown to enhance activation loop phosphorylation ^51^. Intriguingly, however, the arrangement of the kinase domains in both structures is remarkably similar to that observed in DAPK2 and the PKD dimer presented in this study. Due to disorder of the activation loops in both Chk2 and Rad53, which may be the consequence of mutations introduced into the nucleotide binding site or distortions caused by crystal lattice packing, it is impossible to say whether the kinase domains are competent of autophosphorylation in this conformation. Analysis of a spectrum of kinase domain structures, including Chk2 ^51^, has suggested that activation loop exchange may be a common mechanism of *trans*-autophosphorylation ^52, 53^, although these structures also include kinases such as DAPK3 that do not autophosphorylate their activation loop. Activation segment exchange is characterized by a domain swap of the C-terminal portion of the activation loop, which then packs in an equivalent manner against the C-lobe of an opposing protomer. This has been observed in many kinase domain crystal structures ^51, 52, 54–59^, irrespective of whether they autophosphorylate or not. The arrangement of the kinase domains with respect to each other in these structures is variable and the domain swap may reflect energetically favorable lattice packing interactions. Further work will undoubtedly be required to reconcile the differences between dimerization-driven DAPK and PKD autoinhibition and dimerization-mediated activation of Chk2, though it is eminently conceivable that they are simply variations on the same theme and accomplished via similar mechanisms.

The consequences of activation loop phosphorylation in PKD are two-fold: prohibition of inhibitory dimerization and increased catalytic activity. The inhibitory effect of phosphorylation on dimerization has also been reported for Chk2 ^21^. Despite classification as a non-RD kinase on account of a non-canonical HCD motif in its catalytic loop, the catalytic activity of PKD is increased by activation loop phosphorylation. R732 in the activation loop of PKD fulfills the equivalent role of the HRD motif in RD kinases. The inhibitory effect of activation loop phosphorylation on PKD kinase domain dimerization is equally important as the increase in its catalytic activity. Expression of PKD that cannot be phosphorylated in its activation loop leads to the same dominant negative effect on secretion as expression of kinase-dead PKD, despite having considerable basal activity in vitro. Mutation of R732 that reduces, but does not abrogate activation loop phosphorylation in cells and specific activity in vitro, in contrast, does not elicit such a dominant negative effect. PKD signaling therefore depends critically on the acquisition of S742 phosphorylation to prevent autoinhibition. The embryonic lethal phenotype of homozygous S744A/S748A PKD1 knock-in mice ^60^ may reflect the inability of DAG to relieve *trans*-autoinhibition of PKD1 via activation loop autophosphorylation upon binding to the TGN.

The relief of dimerization-mediated autoinhibition of PKD presumably relies on conformational changes in its regulatory domains upon membrane binding. However, in the absence of structural information on the architecture of full-length PKD, it is currently not possible to rationalize this mechanistically. Intriguingly, however, BRAF forms a complex with its substrate kinase MEK1 that bears significant similarity to PKD. Recent work has elucidated the mechanism by which Raf is maintained in an autoinhibited conformation in complex with MEK via intramolecular interactions between its cysteine-rich domain (CRD) domain and a dimer of 14-3-3 proteins ^36^. RAF and MEK form a face-to-face dimer mediated by their αG helices in which both kinases adopt inactive conformations. Displacement of the regulatory domains converts RAF into an active back-to-back dimer that is competent for MEK1 phosphorylation. BRAF contains a Ras-binding domain (RBD) closely followed by its CRD, the combination of which is structurally homologous to the ULD-C1 module of PKD. While the significance of these similarities in domain arrangements is unclear, the activation of BRAF at the membrane by relief of autoinhibition and subsequent acquisition of an active conformation is striking.

Finally, what about PKD inactivation? We have shown here that activation loop phosphorylation prohibits kinase domain dimerization, which implies that its dephosphorylation would be required in order for it to return to its autoinhibited conformation. The identity of the phosphatase that inactivates PKD, however, is not known. Future work will be required to integrate the inactivation of PKD by dephosphorylation into the model of PKD regulation in cells.

## Acknowledgements

We would like to thank Dorothea Anrather, WeiQiang Chen and Markus Hartl in the Max Perutz Labs Mass Spectrometry Facility for help with the acquisition of mass spectra using instruments of the Vienna BioCenter Core Facilities (VBCF). This work was supported by FWF grants P30584 and P33066 to T.A.L. Work in the lab of A.H. is supported by grants from the German Research Foundation (DFG HA357/11-2) and German cancer aid (Grant 70111941). J.E.B. was supported by a Michael Smith Foundation for Health Research (MSFHR) Scholar award (17686) and an operating grant from the Cancer Research Society (CRS-843232).

## Author contributions

R.R. purified all recombinant proteins and performed all the *in vitro* biochemical experiments reported in this study, as well as the cellular membrane translocation assays. T.A.L. performed the *in silico* modeling. M.A.H.P. performed and analyzed the HDX-MS experiments. K.H. performed the FRAP experiments as well as PAUF secretion assays with PKD1 D665N. G.L. performed quantitative Western blotting on cell lysates and PAUF secretion assays with mutant PKD1 variants. S.A.E. performed the Golgi activity reporter assay and analyzed the data. T.H. performed fluorescence microscopy of PKD1 D665N in cells. A.H., J.E.B. and T.A.L. obtained the funding to support the work.

## Materials and Methods

### Reagents, antibodies and plasmids

The plasmid encoding G-PKDrep ^28^ and the plasmids encoding GFP-tagged wild-type PKD1, PKD1 K612W, PKD1 S738/S742E and PKD1 S738/S742E ^61^ were described previously. R732M and D665N mutations were generated by site-directed mutagenesis. The plasmid encoding PAUF-myc/his was kindly provided by Vivek Malhotra (CRG, Barcelona, Spain). The pSer294-specific rabbit polyclonal antibody used for detection of G-PKDrep phosphorylation has been previously described ^28^. Commercially available antibodies used were as follows: anti-phospho-PKD (Ser744/748) rabbit polyclonal antibody, anti-phospho PKD (Ser 916) rabbit polyclonal antibody, anti-myc mouse monoclonal antibody clone 9B11 (all from Cell Signaling Technologies), anti-GFP mouse monoclonal antibody (Roche Diagnostics), anti-tubulin α mouse monoclonal antibody (Merck Chemicals GmbH), and anti-p230 mouse monoclonal antibody (BD Biosciences). Secondary antibodies used were Alexa546 or Alexa488 coupled goat anti–mouse immunoglobulin G (IgG) (Life Technologies), and horseradish peroxidase (HRP) coupled goat anti–mouse and anti–rabbit IgG (Dianova). Nocodazole and doxycyline were purchased from Sigma-Aldrich.

### Protein expression and purification

TEV-cleavable N-terminal GST fusion constructs of PKD1^KD^ (570-892), PKD3^KD^ (563-890), PKD1^ULD-5G-KD^ (48-143-(Gly)_5_-570-892), PKD1^ULD-10G-KD^ (48-143-(Gly)_10_-570-892), and short PKD1^KD^ (570-874) were cloned into the pFastBac Dual vector for expression in Sf9 insect cells. Mutant constructs of PKD1^KD^ were generated by site directed mutagenesis.

Sf9 cell pellets from 1L culture were lysed in 100 mL lysis buffer (50 mM Tris pH 7.5, 150 mM KCl, 1 mM TCEP, 1 mM EDTA, 1 mM EGTA, 0.25 % CHAPS, 20 mM benzamidine, 1 mM PMSF, 4mM MgCl_2_, 1U benzonase, 1x protease inhibitor cocktail (Sigma P8849)). After centrifugation (38 397 g, 4°C, 30 min) the cleared cell lysate was incubated with approximately 3 mL glutathione sepharose beads (Cytiva) for 2 h at 4°C. The beads were then washed with buffer A (50 mM Tris pH 7.5, 150 mM KCl, 1 mM TCEP, 1 mM EDTA, 1 mM EGTA, 0.25 % CHAPS) and dephosphorylated over night at 4°C with 2.2 nmol lambda-phosphatase (purified in house) in 4 mM MnCl_2_. The beads were washed multiple times in buffer A with varying salt concentrations (2x 400mM KCl, 2x 40mM, 2x 150mM), before cleavage with 8.8 nmol TEV protease (purified in house) for 2h at room temperature. Cleaved PKD constructs in solution were separated from the beads, diluted to 37.5 mM KCl in buffer Q_A_ (50 mM Tris pH 8.5, 1 mM TCEP, 1 mM EDTA, 1 % glycerol) and bound to a HiTrap Q HP anion exchange column (Cytiva). After gradient elution with buffer Q_B_ (50 mM Tris pH 8.5, 1 M KCl 1 mM TCEP, 1 mM EDTA, 1 % glycerol) the peak fractions were pooled, concentrated, and injected on a S200 increase 10/300 GL size exclusion column (Cytiva) that was equilibrated in 20 mM Tris pH 7.5, 150 mM KCl, 1 mM TCEP, 1 mM EDTA, 1 % (v/v) glycerol. Constructs with mutated activation loop phosphorylation sites were not treated with lambda phosphatase.

Stoichiometric phosphorylation of PKD1^KD^ (1P) was achieved by incubating the washed, GST-PKD1^KD^ bound beads with 1mM ATP and 5 mM MgCl_2_ at 4°C overnight followed by 1 h at room temperature. After TEV cleavage, the phosphorylated species were separated on a MonoQ 5/50 GL column (Cytiva) and characterized by intact mass spectrometry.

PKD1^ULD^ proteins were purified according to ^14^.

### Analytical size exclusion chromatography

A Superdex 200 increase 3.2/300 size exclusion column (Cytiva), connected to an Äkta Pure with a Micro configuration (Cytiva) was equilibrated in buffer (20 mM HEPES pH 7.4, 150 mM KCl, 1 mM EDTA, 1 mM TCEP, 1 % (v/v) glycerol) at 4°C. 25 µL protein samples were clarified by centrifugation (5 min, 21 000 g) prior to injection at constant flow rate. Peak elution volumes are reported, while peak concentration was determined by integrating the peak area for 20 µL centered on the highest point.

### Differential scanning fluorimetry (DSF)

Thermal stability measurements of PKD1^KD^ were performed in 20 mM Tris pH 7.5, 150 mM KCl, 1 mM TCEP, 1 mM EDTA, 1% (v/v) glycerol with 5x SYPRO Orange (life technologies) using a BioRad iQ™5 Multicolor Real-Time PCR Detection System. A nucleotide-free control was compared to samples containing 5 mM MgCl_2_ and 1 mM ATP or ADP. Each condition was measured at four protein concentrations (1.25, 2.5, 5, 10 µM) and, in the absence of a concentration-dependent change in melting temperature, the measurements were averaged and presented as replicates.

### Cross-linking coupled to mass spectrometry (XL-MS)

#### In-solution cross-linking

Cross-linking of PKD1^KD^ (0P) was performed in 40 mM MES pH 6.5, 100 mM NaCl. 4 µM protein containing 1 mM ATP and 2 mM MgCl_2_ was mixed with varying concentrations of EDC and Sulfo-NHS at a constant ratio of 1:2.5 and incubated for 30 min in the dark. The reaction was quenched with 20 mM β-mercaptoethanol and 50mM Tris pH 7.5. Cross-linked monomeric and dimeric species were separated by SDS-PAGE.

#### Sample preparation for mass spectrometry

The Coomassie-stained gel band was de-stained with a mixture of acetonitrile (Chromasolv®, Sigma-Aldrich) and 50 mM ammonium bicarbonate (Sigma-Aldrich). The proteins were reduced using 10 mM dithiothreitol (Roche) and alkylated with 50 mM iodoacetamide. Trypsin (Promega; Trypsin Gold, Mass Spectrometry Grade) was used for proteolytic cleavage. Digestion was carried out with trypsin at 37°C overnight. Formic acid was used to stop the digestion and the extracted peptides were desalted using C18 Stagetips ^62^.

### Mass spectrometry and data analysis

Peptides were analyzed on an UltiMate 3000 HPLC RSLC nanosystem (Thermo Fisher Scientific) coupled to a Q Exactive HF-X, equipped with a nano-spray ion source using coated emitter tips (PepSep, MSWil). Samples were loaded on a trap column (Thermo Fisher Scientific, PepMap C18, 5 mm × 300 μm ID, 5 μm particles, 100 Å pore size) at a flow rate of 25 μL min^-1^ using 0.1% TFA as mobile phase. After 10 min, the trap column was switched in-line with the analytical C18 column (Thermo Fisher Scientific, PepMap C18, 500 mm × 75 μm ID, 2 μm, 100 Å) and peptides were eluted applying a segmented linear gradient from 2% to 80% solvent B (80% acetonitrile, 0.1% formic acid; solvent A 0.1% formic acid) at a flow rate of 230 nL/min over 120 min. The mass spectrometer was operated in data-dependent mode, survey scans were obtained in a mass range of 350-1600 m/z with lock mass activated, at a resolution of 120,000 at 200 m/z and an AGC target value of 1E6. The 15 most intense ions were selected with an isolation width of 1.2 Thomson for a max. of 150 ms, fragmented in the HCD cell at stepped normalized collision energy at 26%, 28%, and 30%. The spectra were recorded at an AGC target value of 1E5 and a resolution of 60,000. Peptides with a charge of +1, +2, or >+7 were excluded from fragmentation, the peptide match feature was set to preferred, the exclude isotope feature was enabled, and selected precursors were dynamically excluded from repeated sampling for 20 seconds within a mass tolerance of 8 ppm.

Raw data were processed using the MaxQuant software package 1.6.17.0 ^63^ and searched against the Uniprot human reference proteome (January 2020, www.uniprot.org) as well as a database of most common contaminants. The search was performed with standard identification settings: full trypsin specificity allowing a maximum of two missed cleavages. Carbamidomethylation of cysteine residues was set as fixed, oxidation of methionine and acetylation of protein N-termini as variable modifications. All other settings were left at default. Results were filtered at a false discovery rate of 1% at protein and peptide spectrum match level. To identify cross-linked peptides, the spectra were searched using pLink software 2.3.9 ^64^ against the sequences of the top 8 non-contaminant proteins from the MQ search sorted by iBAQ. Carbamidomethylation of cysteine was set as fixed, oxidation of methionine and acetylation of protein N-termini as variable modifications. The enzyme specificity was set to trypsin allowing 4 missed cleavage sites. Crosslinker settings were selected as EDC. Search results were filtered for 1% FDR (false discovery rate) on the PSM level (peptide-spectrum matches) and a maximum precursor mass deviation of 5 ppm. To remove low quality PSMs, additionally an e-Value cutoff of < 0.001 was applied. Cross-link maps were generated in xiNET ^65^.

### Proteomics data deposition

The mass spectrometry proteomics data have been deposited to the ProteomeXchange Consortium via the PRIDE partner repository ^66^ with the dataset identifier PXD031997.

### *In silico* modeling

Homology modeling of the PKD1 kinase domain (residues 576-873) was performed using the Rosetta Comparative Modeling tool ^19^ and the structures of phosphorylase kinase (PDB 2phk), Chk2 (PDB 3i6u), DAPK2 (PDB 2a2a; 2yaa) as templates. The active state was modeled using ligand coordinate and parameter files for ATP calculated for ATP from the active conformation of phosphorylase kinase (PDB 2phk) and the built-in Rosetta parameter file for magnesium. The model was compared to the AlphaFold2 ^20^ prediction for the PKD1 kinase domain and tested biochemically.

Homology modeling of the PKD1 kinase domain dimer was performed using the Rosetta Comparative Modeling tool and symmetry imposed by the structure of Chk2 (PDB 3i6u). Additional constraints were imposed based on dimer-specific intramolecular EDC cross-links (615-622) and intermolecular EDC cross-links (622-831), and an EDC cross-link common to both monomer and dimer (616-624), with a distance constraint of 3.8 Å. The model was tested biochemically by introducing a single point mutation (E787K) into the predicted dimer interface.

### Size exclusion chromatography coupled to multi-angle light scattering (SEC-MALS)

For determination of particle mass and polydispersity, 60 µL of purified PKD1^KD^ at 2 mg/mL was injected onto an S200 10/300 column (Cytiva) connected to a 1260 Infinity HPLC (Agilent Technologies). A MiniDawn Treos (Wyatt) was used to detect light scattering at 690 nm and a Shodex RI-101 (Shodex) detector was used for refractive index measurement. All runs were performed at room temperature in 20 mM HEPES pH 7.4, 150 mM KCl, 1 mM EDTA, 1 mM TCEP, 1 % (v/v) glycerol.

### Kinase assays

All kinase assays were done with radiolabeled [γ-32P] ATP (Hartman Analytic) in a reaction buffer containing 50mM HEPES pH 7.4, 150 mM KCl, 1 mM TCEP, 1 mM EDTA, 1 % (v/v) glycerol, 0.05 % (v/v) Tween20, 1 mM ATP, 5 mM MgCl_2_, 1 µL [γ-32P] ATP /100 µL reaction and were terminated by the addition of 10 mM EDTA. The radioactivity from phosphorylated material immobilized in polyacrylamide gels or on 0.45 μm nitrocellulose membranes (Cytiva) was detected by exposure to a phosphor screen, imaged by an Amersham Typhoon phosphorimager, quantified in ImageJ, and converted into molar quantities by calibration with internal standards.

Assays probing the fraction of autophosphorylated PKD^KD^ at the reaction plateau at different concentrations were performed at 30°C overnight. After termination of the reaction, aliquots corresponding to 0.5 pmol of PKD^KD^ were subjected to SDS-PAGE to separate the phosphorylated protein from excess [γ-32P] ATP. The gels were stained with silver stain (Invitrogen), dried and exposed to a phosphor screen overnight. The fraction of phosphorylated PKD^KD^ was calculated from the internal standard, assuming a single phosphate is incorporated per PKD^KD^ molecule (confirmed by intact mass spectrometry).

For autophosphorylation assays probing the *trans*-autoinhibitory potential of PKD1^KD^ (D706N/2E), 50 nM of active PKD1^KD^ (0P) or PKD1^KD^ (E787K) was mixed with 0.5 µM or 5 µM of kinase dead PKD1^KD^ (D706N/2E), and incubated over night at 30°C. Aliquots corresponding to 0.5 pmol of active kinase were subjected to SDS-PAGE. The gels were washed three times in dH_2_O and processed as described above. The signal was normalized to PKD1^KD^ (0P) at 50 nM.

Autophosphorylation time course experiments were performed at 30°C with 50 nM PKD1^KD^. Samples were taken from the reaction volume, terminated and aliquots corresponding to 0.5 pmol of PKD1^KD^ were subjected to SDS-PAGE and processed as described above.

Substrate phosphorylation time course assays with PKD1^ULD-5G/10G-KD^ were performed on ice with 50 nM protein and 100 µM biotinylated Syntide2 (biotin-PLARTSVAG (GenScript)). Samples were taken from the reaction volume, terminated, and spotted on a nitrocellulose membrane. The membrane was washed five times with 20 ml of 75 mM H_3_PO_4_ and exposed to a phosphor screen for 2 h together with a set of internal standards.

Michaelis-Menten kinetics were performed on ice with 0-150 µM biotin-Syntide 2. The reactions were started by adding 10 nM PKD1^KD^ and terminated after 2 min. Aliquots were spotted onto a nitrocellulose membrane and processed as described above.

Substrate phosphorylation assays with PKD1^KD^ (D665N) were done with 50 nM PKD1^KD^ (D665N) and 0-100 µM Syntide 2 (Sigma).

For the analysis of PKD1 autophosphorylation by intact mass spectrometry 20 mL of kinase reaction (20 mM HEPES pH 7.4, 150 mM KCl, 1 mM TCEP, 1 mM EDTA, 1 % glycerol, 0.25 % CHAPS, 5 mM MgCl_2_ 1 mM ATP, 10 nM short PKD1^KD^, 40 nM PKD1^KD^ D706N) were incubated at 30°C for 7 h. The solution was concentrated to 70 µL in a 10 kDa cutoff Vivaspin concentrator (Sartorius) and the protein was buffer exchanged with 20 mM HEPES pH 7.4, 150 mM KCl, 1 mM TCEP, 1 mM EDTA, 1 % glycerol in a Zeba desalting column (Thermo Fisher Scientific). An unphosphorylated reference sample was prepared with 2 µM short PKD1^KD^ and 8 µM PKD1^KD^ D706N in the same buffer.

### HDX-MS

#### Sample preparation

HDX reactions for wild-type PKD1^KD^ and PKD1^KD^ S738/742E were conducted in a final reaction volume of 6 µL with a final concentration of 5 µM (30 pmol) PKD1^KD^ or PKD1^KD^ S738/742E. The reaction was initiated by the addition of 5.25 µL of D_2_O buffer (20 mM pH 7.5 HEPES, 100mM NaCl, 1 mM ATPγS, 5 mM MgCl_2_, 85.8% D2O (V/V)) to 0.75 µL of PKD1cat WT or mutant (final D_2_O concentration of 71.3%). The reaction proceeded for 3, 30, 300, or 3000s at 20°C, before being quenched with ice cold acidic quench buffer, resulting in a final concentration of 0.6M guanidine-HCl and 0.9% formic acid post quench. All conditions and timepoints were generated in triplicate. Samples were flash frozen immediately after quenching and stored at -80°C until injected onto the ultra-performance liquid chromatography (UPLC) system for proteolytic cleavage, peptide separation, and injection onto a QTOF for mass analysis, described below.

#### Protein digestion and MS/MS data collection

Protein samples were rapidly thawed and injected onto an integrated fluidics system containing a HDx-3 PAL liquid handling robot and climate-controlled (2°C) chromatography system (LEAP Technologies), a Dionex Ultimate 3000 UHPLC system, as well as an Impact HD QTOF Mass spectrometer (Bruker). The full details of the automated LC system are described in ^67^. The protein was run over one immobilized pepsin column (Trajan; ProDx protease column, 2.1 mm x 30 mm PDX.PP01-F32) at 200 µL/min for 3 minutes at 8°C. The resulting peptides were collected and desalted on a C18 trap column (Acquity UPLC BEH C18 1.7mm column (2.1 x 5 mm); Waters 186003975). The trap was subsequently eluted in line with an ACQUITY 1.7 μm particle, 100 × 1 mm^2^ C18 UPLC column (Waters), using a gradient of 3-35% B (Buffer A 0.1% formic acid; Buffer B 100% acetonitrile) over 11 minutes immediately followed by a gradient of 35-80% over 5 minutes. Mass spectrometry experiments acquired over a mass range from 150 to 2200 m/z using an electrospray ionization source operated at a temperature of 200°C and a spray voltage of 4.5 kV.

#### Peptide identification

Peptides were identified from the non-deuterated samples of PKD1^KD^ or PKD1^KD^ S738/742E using data-dependent acquisition following tandem MS/MS experiments (0.5 s precursor scan from 150-2000 m/z; twelve 0.25 s fragment scans from 150-2000 m/z). MS/MS datasets were analyzed using PEAKS7 (PEAKS), and peptide identification was carried out by using a false discovery-based approach, with a threshold set to 0.1% using a database of known contaminants found in Sf9 cells ^68^. The search parameters were set with a precursor tolerance of 20 ppm, fragment mass error 0.02 Da, charge states from 1-8, leading to a selection criterion of peptides that had a -10logP score of 26.2 and 24.8 for PKD1^KD^ or PKD1^KD^ S738/742E, respectively.

#### Mass Analysis of Peptide Centroids and Measurement of Deuterium Incorporation

HD-Examiner Software (Sierra Analytics) was used to calculate the level of deuterium incorporation into each peptide. All peptides were manually inspected for correct charge state, correct retention time, and appropriate selection of isotopic distribution. Deuteration levels were calculated using the centroid of the experimental isotope clusters. Results are presented as relative levels of deuterium incorporation, with no correction for back exchange. The only correction was for the deuterium percentage of the buffer in the exchange reaction (71.3%). Differences in exchange in a peptide were considered significant if they met all three of the following criteria: ≥5% change in exchange, ≥0.5 Da difference in exchange, and a 2-tailed T-test value of less than 0.01 at any time point. The raw HDX data are shown in two different formats. The raw peptide deuterium incorporation graphs for a selection of peptides with significant differences are shown in Supplementary Figure S5A, with the raw data for all analyzed peptides in the source data. To allow for visualization of differences across all peptides, we utilized number of deuteron difference (#D) plots (Figure 3B). These plots show the total difference in deuterium incorporation over the entire H/D exchange time course, with each point indicating a single peptide. Samples were only compared within a single experiment and were never compared to experiments completed at a different time with a different final D_2_O level. The data analysis statistics for all HDX-MS experiments are in Supplemental Table 2 according to published guidelines ^69^. The mass spectrometry proteomics data have been deposited to the ProteomeXchange Consortium via the PRIDE partner repository ^66^ with the dataset identifier PXD033139.

### Cell culture and transfection

HeLa and HEK293T cells were maintained in RPMI 1640 medium supplemented with 10% FCS. Cell lines were authenticated using Multiplex Cell Authentication by Multiplexion (Heidelberg, Germany) as described recently (Castro et al., 2013). The SNP profiles matched known profiles or were unique. Cells were tested negative for mycoplasma contamination using MycoAlert (Lonza, Switzerland). For transient plasmid transfections, HEK293T cells were transfected with TransIT-293 (Mirus Bio, Madison, WI, USA). HeLa cells were transfected with TransIT-HeLaMONSTER (Mirus Bio).

Flp-In T-REx-HeLa cells (generated by Elena Dobrikova and Matthias Gromeier, Duke University Medical Center, Durham, NC, USA) were grown in DMEM containing 10% FCS, 100 mg/ml zeocin and 10 mg/ml blasticidin (HeLa). These cells stably express the Tet repressor, contain a single Flp Recombination Target (FRT) site and were used to generate the Flp-In-T-REx-EGFP-GEF-H1-lines. Cells were cotransfected with pcDNA5/FRT/TO-EGFP-GEF-H1 wt or C53R and the Flp recombinase expression plasmid pOG44 at a ratio of 1:10 and then selected with 500 mg/ml hygromycin B. Induction of protein expression with doxycycline was at 10 ng/ml.

### Immunofluorescence staining and confocal microscopy

Cells expressing PKD1-EGFP were grown on glass coverslips coated with 2,5 mg/ml collagen R (Serva, Heidelberg, Germany) and fixed for 15 min with 4% (v/v) paraformaldehyde. After washes in PBS, cells were incubated for 5 min with 1 M glycine in PBS and permeabilized for 2 min with 0.1% (v/v) Triton X-100 in PBS. Blocking was performed with 5% (v/v) bovine serum (PAN) in PBS for 30 min. Fixed cells were incubated with primary antibodies diluted in blocking buffer for 2 hr at room temperature. Following three washing steps with PBS, cells were incubated with Alexa-Fluor-546-labeled secondary antibodies in blocking buffer for 1 hr at room temperature. Nuclei were counterstained with DAPI and mounted in ProLong Gold Antifade Reagent (Thermo Fisher Scientific). All samples were analyzed at room temperature using a confocal laser scanning microscope (LSM 710, Carl Zeiss) equipped with a Plan Apochromat 63x/1.40 DIC M27 (Carl Zeiss, Jena, Germany) oil-immersion objective. GFP was excited with the 488 nm line of an Argon laser, its emission was detected from 496 to 553 nm. Alexa546 was excited with a 561 nm DPSS laser, its emission was detected from 566 to 622 nm. Image acquisition for G-PKDrep ratiometric imaging was done as follows: z-stacks of 0.5 mm intervals were acquired throughout the cell and maximum intensity projections were calculated. GFP and Alexa546 channels were hereby acquired with the same pinhole setting that was adjusted to 1 AU in the Alexa 546 channel. Laser powers were adjusted to prevent fluorophore saturation and identical photomultiplier tube and laser settings were maintained throughout the whole experiment. Image processing and analysis was performed with Zen black 2.1 software. Regions of interest of identical Golgi areas of reporter expressing cells were selected in the GFP channel, mean pixel intensity values of the selected areas in both channels were measured and the Alexa546 to GFP ratio was calculated.

### Protein extraction of cells and Western blotting

Whole cell extracts were obtained by solubilizing cells in lysis buffer (20 mM Tris pH 7.4, 150 mM NaCl, 1% Triton X-100, 1 mM EDTA, 1 mM ethylene glycol tetra acetic acid (EGTA), plus Complete protease inhibitors and PhosSTOP (Roche Diagnostics, Basel, Switzerland)). Whole cell lysates were clarified by centrifugation for 15 min at 16,000 g and 4°C. Equal amounts of protein were loaded on 10% polyacrylamide gels or were run on NuPage Novex 4–12% Bis-Tris gels (Life Technologies) and blotted onto nitrocellulose membranes using the iBlot device (Life Technologies). Membranes were blocked for 30 min with 0.5% (v/v) blocking reagent (Roche Diagnostics) in PBS containing 0.05% (v/v) Tween-20. Membranes were incubated with primary antibodies overnight at 4°C, followed by 1 hr incubation with HRP-conjugated or IRDye-conjugated secondary antibodies at room temperature.

Proteins were visualized using an enhanced chemiluminescence detection system (Thermo Fisher Scientific, Waltham, MA, USA) or the Licor Odyssey system. Special care was taken not to overexpose in order to guarantee accurate quantifications. Densitometry was performed using Image Studio Lite 4.0 (Li-COR Biosciences, Bad Homburg, Germany). For each protein, the integrated density of the signal was measured and corrected for background signals.

### PAUF secretion assay

FlpIN Trex HeLa cells stably expressing PKD1-EGFP variants WT, K612W, and D655N were transfected with a plasmid encoding PAUF-Myc-His and after another 24 hours, the expression of PKD1-EGFP was induced with doxycycline. HEK293T cells were transiently transfected with plasmids encoding PKD1-EGFP variants WT, S2E, S2A, R732M, or R732M S2E and PAUF-Myc-His in a ratio of 1:5. Growth medium was removed 24 h post-transfection, and cells were incubated in serum-free medium. After 5 h, the medium was collected, and cells were lysed. Aliquots of medium and detergent-soluble cell proteins were assayed for PAUF content by western blot analysis.

### FRAP

The diffusion mobility of PKD1-GFP in living HeLa cells was analyzed using a laser scanning confocal microscope. Cells were imaged with a Plan Apochromat 63×/1.4 DIC M27 objective lens. An initial prebleach image was taken, followed by bleaching the Golgi region of interest five times with high-intensity laser light (488 nm line, 80% laser power). Recovery of fluorescence in the bleached area was measured over time by scanning every second. Fluorescence intensity values were normalized to the unbleached region. Fitting of curves was performed with a one-phase exponential equation (Y = Ymax[1 − exp(−K*X)]; Prism 9, GraphPad Software, La Jolla, CA).

### In-cell membrane translocation assay

For transfection into mammalian cells, wild type and mutant constructs of full-length PKD1 were cloned into the EGFP-C1 and mCherry-C1 vectors (Clontech), resulting in N-terminally labelled fusion proteins. With the exception of PKD1^K612N^, which was expressed as an EGFP fusion, all mutant PKD1 variants were expressed as mCherry fusions.

COS7 cells were seeded into a 4-well chamber microscopy dish (InVitro Scientific) and co-transfected with wild type and mutant constructs of PKD 20 h post-seeding with 250 ng of wild-type and 250 ng mutant plasmid DNA per well using TurboFect (Thermo Scientific) according to the manufacturer’s instructions. After 24 h, the medium was exchanged to Hank’s Buffered Saline Solution (Life Technologies), and cells were imaged in HBSS on a Zeiss LSM710 confocal microscope equipped with 488- and 561-nm lasers. Membrane translocation in live cells was induced by the addition of 500 nM phorbol 12-myristate 13-acetate (PMA; Sigma) and monitored by imaging in 20–30 s intervals. The decay of cytosolic fluorescence over time was quantified in a region of interest using ImageJ, and the half-maximum translocation time (*t*_0.5_) was determined by a logistic fit function over the decay curve. Mutations were considered to impact membrane translocation if the ratio of *t*_0.5_ for the wild type over the mutant protein deviated significantly from 1. For visualization, the data curves were normalized to the minimum and maximum values of fluorescence in the cell.

## References

1. Manning, G. The Protein Kinase Complement of the Human Genome. Science (80-.). 298, 1912–1934 (2002).

2. Endicott, J. A., Noble, M. E. M. & Johnson, L. N. The structural basis for control of eukaryotic protein kinases. Annu. Rev. Biochem. 81, 587–613 (2012).

3. Modi, V. & Dunbrack, R. L. Defining a new nomenclature for the structures of active and inactive kinases. Proc. Natl. Acad. Sci. U. S. A. 116, 6818–6827 (2019).

4. Lapierre, J.-M. et al. Discovery of 3-(3-(4-(1-Aminocyclobutyl)phenyl)-5-phenyl-3H-imidazo[4,5-b]pyridin-2-yl)pyridin-2-amine (ARQ 092): An Orally Bioavailable, Selective, and Potent Allosteric AKT Inhibitor. J. Med. Chem. 59, 6455–69 (2016).

5. Noble, M. E. M., Endicott, J. A. & Johnson, L. N. Protein kinase inhibitors: insights into drug design from structure. Science 303, 1800–5 (2004).

6. Nolen, B., Taylor, S. & Ghosh, G. Regulation of protein kinases; controlling activity through activation segment conformation. Mol. Cell 15, 661–75 (2004).

7. Beenstock, J., Mooshayef, N. & Engelberg, D. How Do Protein Kinases Take a Selfie (Autophosphorylate)? Trends Biochem. Sci. 41, 938–953 (2016).

8. Dodson, C. A., Yeoh, S., Haq, T. & Bayliss, R. A kinetic test characterizes kinase intramolecular and intermolecular autophosphorylation mechanisms. Sci. Signal. 6, ra54 (2013).

9. Liljedahl, M. et al. Protein kinase D regulates the fission of cell surface destined transport carriers from the trans-Golgi network. Cell 104, 409–20 (2001).

10. Wakana, Y. et al. A new class of carriers that transport selective cargo from the trans Golgi network to the cell surface. EMBO J. 31, 3976–3990 (2012).

11. Van Lint, J. V, Sinnett-Smith, J. & Rozengurt, E. Expression and characterization of PKD, a phorbol ester and diacylglycerol-stimulated serine protein kinase. J. Biol. Chem. 270, 1455–61 (1995).

12. Baron, C. L. & Malhotra, V. Role of diacylglycerol in PKD recruitment to the TGN and protein transport to the plasma membrane. Science 295, 325–8 (2002).

13. Rybin, V. O., Guo, J. & Steinberg, S. F. Protein kinase D1 autophosphorylation via distinct mechanisms at Ser744/Ser748 and Ser916. J. Biol. Chem. 284, 2332–43 (2009).

14. Elsner, D. J., Siess, K. M., Gossenreiter, T., Hartl, M. & Leonard, T. A. A ubiquitin-like domain controls protein kinase D dimerization and activation by trans-autophosphorylation. J. Biol. Chem. 294, 14422–14441 (2019).

15. Waldron, R. T. et al. Activation loop Ser744 and Ser748 in protein kinase D are transphosphorylated in vivo. J. Biol. Chem. 276, 32606–15 (2001).

16. Hausser, A. et al. Protein kinase D regulates vesicular transport by phosphorylating and activating phosphatidylinositol-4 kinase IIIbeta at the Golgi complex. Nat. Cell Biol. 7, 880–6 (2005).

17. Fugmann, T. et al. Regulation of secretory transport by protein kinase D-mediated phosphorylation of the ceramide transfer protein. J. Cell Biol. 178, 15–22 (2007).

18. Nhek, S. et al. Regulation of oxysterol-binding protein Golgi localization through protein kinase D-mediated phosphorylation. Mol. Biol. Cell 21, 2327–37 (2010).

19. Song, Y. et al. High-resolution comparative modeling with RosettaCM. Structure 21, 1735–42 (2013).

20. Jumper, J. et al. Highly accurate protein structure prediction with AlphaFold. Nature 1–7 (2021) doi:10.1038/s41586-021-03819-2.

21. Cai, Z., Chehab, N. H. & Pavletich, N. P. Structure and activation mechanism of the CHK2 DNA damage checkpoint kinase. Mol. Cell 35, 818–29 (2009).

22. DiMaio, F., Leaver-Fay, A., Bradley, P., Baker, D. & André, I. Modeling Symmetric Macromolecular Structures in Rosetta3. PLoS One 6, e20450 (2011).

23. Iglesias, T., Waldron, R. T. & Rozengurt, E. Identification of in vivo phosphorylation sites required for protein kinase D activation. J. Biol. Chem. 273, 27662–7 (1998).

24. Reinhardt, R., Truebestein, L., Schmidt, H. A. & Leonard, T. A. It Takes Two to Tango: Activation of Protein Kinase D by Dimerization. Bioessays e1900222 (2020) doi:10.1002/bies.201900222.

25. Brinkworth, R. I., Breinl, R. A. & Kobe, B. Structural basis and prediction of substrate specificity in protein serine/threonine kinases. Proc. Natl. Acad. Sci. U. S. A. 100, 74–79 (2003).

26. Creixell, P. et al. Kinome-wide decoding of network-attacking mutations rewiring cancer signaling. Cell 163, 202–17 (2015).

27. Fuchs, Y. F. et al. A Golgi PKD activity reporter reveals a crucial role of PKD in nocodazole-induced Golgi dispersal. Traffic 10, 858–67 (2009).

28. Fuchs, Y. F. et al. A Golgi PKD activity reporter reveals a crucial role of PKD in nocodazole-induced Golgi dispersal. Traffic 10, 858–67 (2009).

29. Lučić, I., Truebestein, L. & Leonard, T. A. Novel Features of DAG-Activated PKC Isozymes Reveal a Conserved 3-D Architecture. J. Mol. Biol. 428, 121–141 (2016).

30. Hein, M. Y. et al. A human interactome in three quantitative dimensions organized by stoichiometries and abundances. Cell 163, 712–23 (2015).

31. Truebestein, L. et al. Structure of autoinhibited Akt1 reveals mechanism of PIP3- mediated activation. Proc. Natl. Acad. Sci. U. S. A. 118, (2021).

32. Kim, C., Cheng, C. Y., Saldanha, S. A. & Taylor, S. S. PKA-I Holoenzyme Structure Reveals a Mechanism for cAMP-Dependent Activation. Cell 130, 1032–1043 (2007).

33. Dar, A. C., Dever, T. E. & Sicheri, F. Higher-order substrate recognition of eIF2alpha by the RNA-dependent protein kinase PKR. Cell 122, 887–900 (2005).

34. Song, H. et al. Phosphoprotein–Protein Interactions Revealed by the Crystal Structure of Kinase-Associated Phosphatase in Complex with PhosphoCDK2. Mol. Cell 7, 615–626 (2001).

35. Haling, J. R. et al. Structure of the BRAF-MEK complex reveals a kinase activity independent role for BRAF in MAPK signaling. Cancer Cell 26, 402–413 (2014).

36. Park, E. et al. Architecture of autoinhibited and active BRAF-MEK1-14-3-3 complexes. Nature 1–5 (2019) doi:10.1038/s41586-019-1660-y.

37. Levina, A., Fleming, K. D., Burke, J. E. & Leonard, T. A. Activation of the essential kinase PDK1 by phosphoinositide-driven trans-autophosphorylation. Nat. Commun. 13, 1874 (2022).

38. Simon, B. et al. Death-Associated Protein Kinase Activity Is Regulated by Coupled Calcium/Calmodulin Binding to Two Distinct Sites. Structure 24, 851–61 (2016).

39. Patel, A. K. et al. Structure of the Dimeric Autoinhibited Conformation of DAPK2, a Pro-Apoptotic Protein Kinase. J. Mol. Biol 409, 369–383 (2011).

40. Graves, P. R., Winkfield, K. M. & Haystead, T. A. J. Regulation of zipper-interacting protein kinase activity in vitro and in vivo by multisite phosphorylation. J. Biol. Chem. 280, 9363–74 (2005).

41. Tereshko, V., Teplova, M., Brunzelle, J., Watterson, D. M. & Egli, M. Crystal structures of the catalytic domain of human protein kinase associated with apoptosis and tumor suppression. Nat. Struct. Biol. 8, 899–907 (2001).

42. Lochhead, P. A. Protein kinase activation loop autophosphorylation in cis: overcoming a Catch-22 situation. Sci. Signal. 2, pe4 (2009).

43. Griffith, J. et al. The structural basis for autoinhibition of FLT3 by the juxtamembrane domain. Mol. Cell 13, 169–78 (2004).

44. Hubbard, S. R., Wei, L., Ellis, L. & Hendrickson, W. A. Crystal structure of the tyrosine kinase domain of the human insulin receptor. Nature 372, 746–54 (1994).

45. Hubbard, S. R. Crystal structure of the activated insulin receptor tyrosine kinase in complex with peptide substrate and ATP analog. EMBO J. 16, 5572–81 (1997).

46. Matsuoka, S. et al. Ataxia telangiectasia-mutated phosphorylates Chk2 in vivo and in vitro. Proc. Natl. Acad. Sci. U. S. A. 97, 10389–94 (2000).

47. Melchionna, R., Chen, X. B., Blasina, A. & McGowan, C. H. Threonine 68 is required for radiation-induced phosphorylation and activation of Cds1. Nat. Cell Biol. 2, 762–5 (2000).

48. Ahn, J.-Y., Li, X., Davis, H. L. & Canman, C. E. Phosphorylation of threonine 68 promotes oligomerization and autophosphorylation of the Chk2 protein kinase via the forkhead-associated domain. J. Biol. Chem. 277, 19389–95 (2002).

49. Li, J. et al. Chk2 oligomerization studied by phosphopeptide ligation: implications for regulation and phosphodependent interactions. J. Biol. Chem. 283, 36019–30 (2008).

50. Wybenga-Groot, L. E. et al. Structural basis of Rad53 kinase activation by dimerization and activation segment exchange. Cell. Signal. 26, 1825–36 (2014).

51. Oliver, A. W. et al. Trans-activation of the DNA-damage signalling protein kinase Chk2 by T-loop exchange. EMBO J. 25, 3179–90 (2006).

52. Pike, A. C. W. et al. Activation segment dimerization: a mechanism for kinase autophosphorylation of non-consensus sites. EMBO J. 27, 704–14 (2008).

53. Oliver, A. W., Knapp, S. & Pearl, L. H. Activation segment exchange: a common mechanism of kinase autophosphorylation? Trends Biochem. Sci. 32, 351–356 (2007).

54. Lawhorn, B. G. et al. Identification of Purines and 7-Deazapurines as Potent and Selective Type I Inhibitors of Troponin I-Interacting Kinase (TNNI3K). J. Med. Chem. 58, 7431–7448 (2015).

55. Taylor, C. A. et al. Domain-Swapping Switch Point in Ste20 Protein Kinase SPAK. Biochemistry 54, 5063–71 (2015).

56. Marcotte, D. et al. Germinal-center kinase-like kinase co-crystal structure reveals a swapped activation loop and C-terminal extension. Protein Sci. 26, 152–162 (2017).

57. Mayo, C. B. et al. Structural Basis of Protein Kinase R Autophosphorylation. Biochemistry 58, 2967–2977 (2019).

58. Wu, P. et al. Hematopoietic Progenitor Kinase-1 Structure in a Domain-Swapped Dimer. Structure 27, 125–133.e4 (2019).

59. Lim, D. C. et al. Redox priming promotes Aurora A activation during mitosis. Sci. Signal. 13, (2020).

60. Matthews, S. A. et al. Unique functions for protein kinase D1 and protein kinase D2 in mammalian cells. Biochem. J. 432, 153–63 (2010).

61. Hausser, A. et al. Structural requirements for localization and activation of protein kinase C μ (PKCμ) at the Golgi compartment. J. Cell Biol. 156, 65–74 (2002).

62. Rappsilber, J., Mann, M. & Ishihama, Y. Protocol for micro-purification, enrichment, pre-fractionation and storage of peptides for proteomics using StageTips. Nat. Protoc. 2, 1896–906 (2007).

63. Tyanova, S., Temu, T. & Cox, J. The MaxQuant computational platform for mass spectrometry-based shotgun proteomics. Nat. Protoc. 11, 2301–2319 (2016).

64. Chen, Z. L. et al. A high-speed search engine pLink 2 with systematic evaluation for proteome-scale identification of cross-linked peptides. Nat. Commun. 2019 101 10, 1–12 (2019).

65. Combe, C. W., Fischer, L. & Rappsilber, J. xiNET: Cross-link Network Maps With Residue Resolution. Mol. Cell. Proteomics 14, 1137–1147 (2015).

66. Perez-Riverol, Y. et al. The PRIDE database resources in 2022: a hub for mass spectrometry-based proteomics evidences. Nucleic Acids Res. 50, D543–D552 (2022).

67. Stariha, J. T. B., Hoffmann, R. M., Hamelin, D. J. & Burke, J. E. Probing Protein-Membrane Interactions and Dynamics Using Hydrogen-Deuterium Exchange Mass Spectrometry (HDX-MS). Methods Mol. Biol. 2263, 465–485 (2021).

68. Dobbs, J. M., Jenkins, M. L. & Burke, J. E. Escherichia coli and Sf9 Contaminant Databases to Increase Efficiency of Tandem Mass Spectrometry Peptide Identification in Structural Mass Spectrometry Experiments. J. Am. Soc. Mass Spectrom. 31, 2202–2209 (2020).

69. Masson, G. R. et al. Recommendations for performing, interpreting and reporting hydrogen deuterium exchange mass spectrometry (HDX-MS) experiments. Nat. Methods 16, 595–602 (2019).

