## Supplementary Information for "PKD autoinhibition in *trans* regulates activation loop autophosphorylation in *cis*"

Supplementary Figures S1-S9
Supplementary Tables S1-S2

**Figure S1. Mass spectra of all protein constructs employed in this study.**

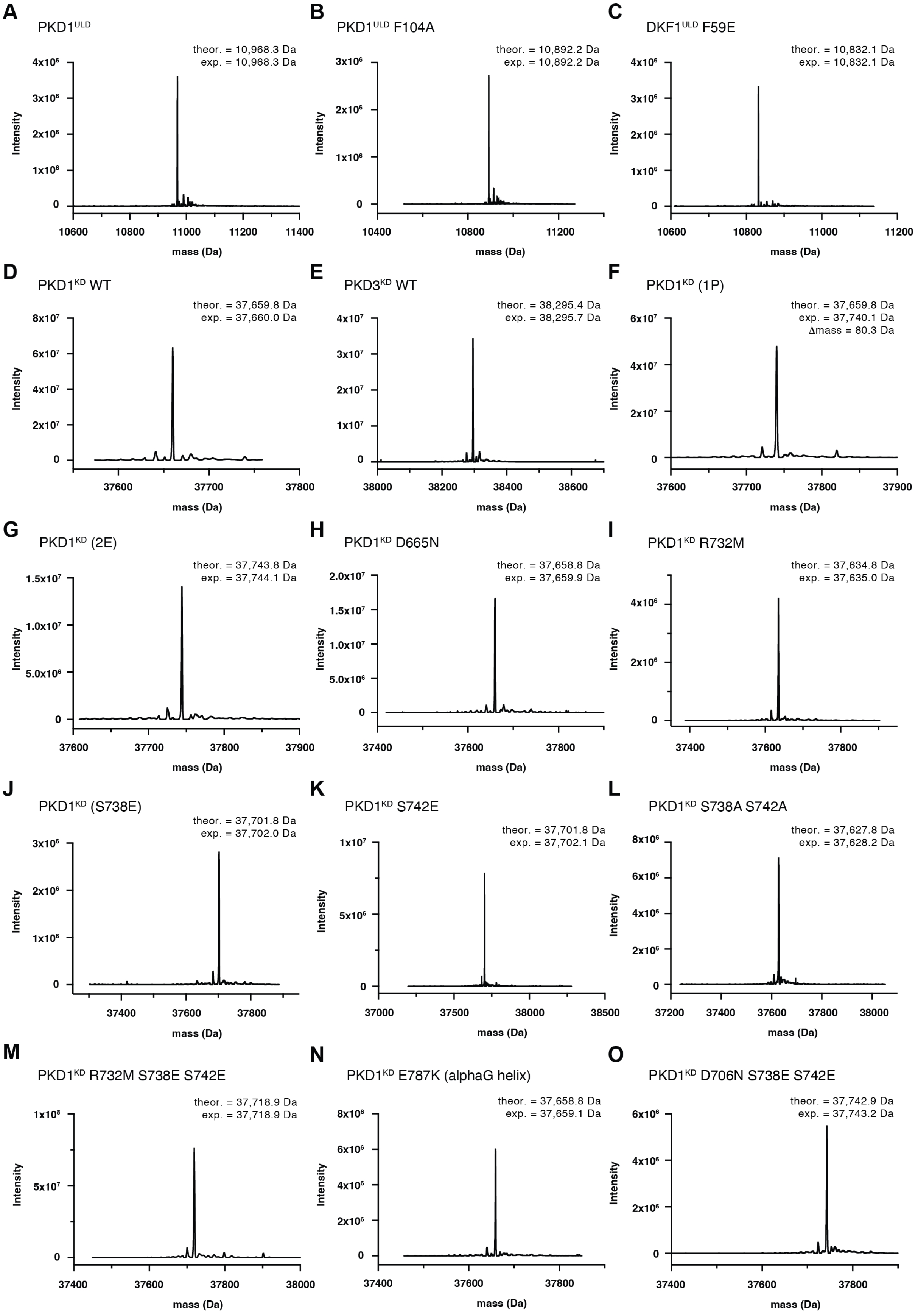

**Figure S2. Mass spectra of all protein constructs employed in this study.**

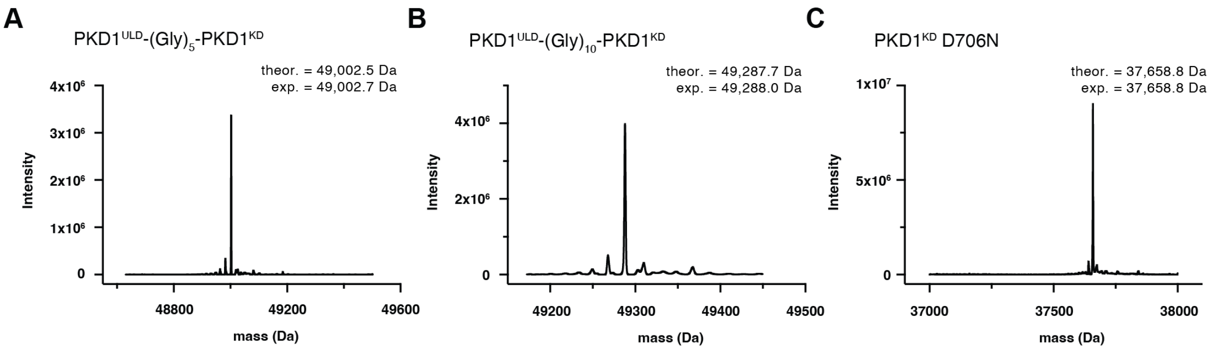

**Figure S3. The PKD kinase domain forms a face-to-face dimer.**

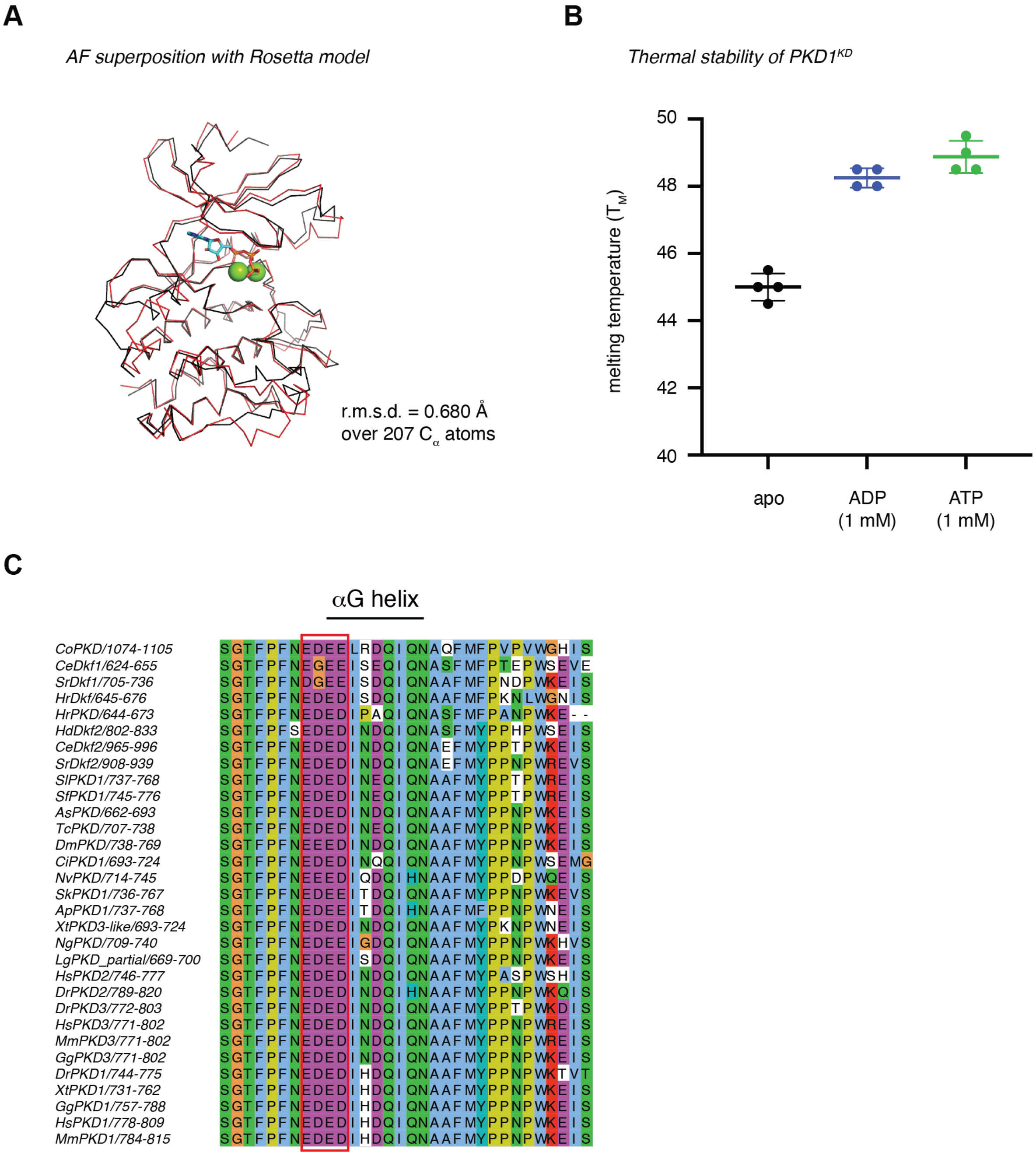


A. Superposition of *in silico* models of PKD1 kinase domain derived from Rosetta and AlphaFold.

B. Thermal stability of PKD1$^{KD}$ in the absence (black) or presence of 1 mM ADP (blue) or ATP (green).

C. Sequence conservation of the αG helix of PKD orthologs.

**Figure S4. Dimerization of the PKD kinase domain is autoinhibitory.**

A B

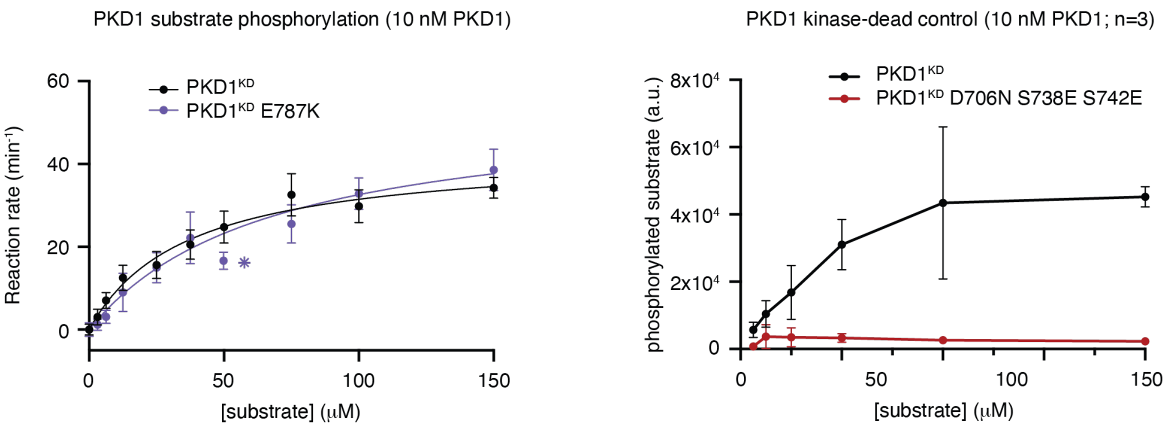


| | n | $K_M$ (μM) | $V_{max}$ (pmol/min) | $V_{max}$ fold change |
|---|---|---|---|---|
| PKD1KD | 10 | 36.4 ± 6.4 | 42.9 ± 2.8 | - |
| PKD1KD E787K | 4 | 67.4 ± 13.6 | 54.5 ± 5.0 | 1.3 |

A. Substrate phosphorylation kinetics of PKD1KD E787K. Error bars are the standard deviation of *n* biologically independent experiments. Table indicates the number (*n*) of independent biological replicates and the values for $K_M$ and $V_{max}$ derived from fitting the data with the Michaelis-Menten equation. Asterisk denotes removal of one outlier.

B. Substrate phosphorylation kinetics of PKD1KD D706N S738E S742E. Error bars are the standard deviation of 3 biologically independent experiments.

**Figure S5. Activation loop autophosphorylation increases PKD1 catalytic activity.**

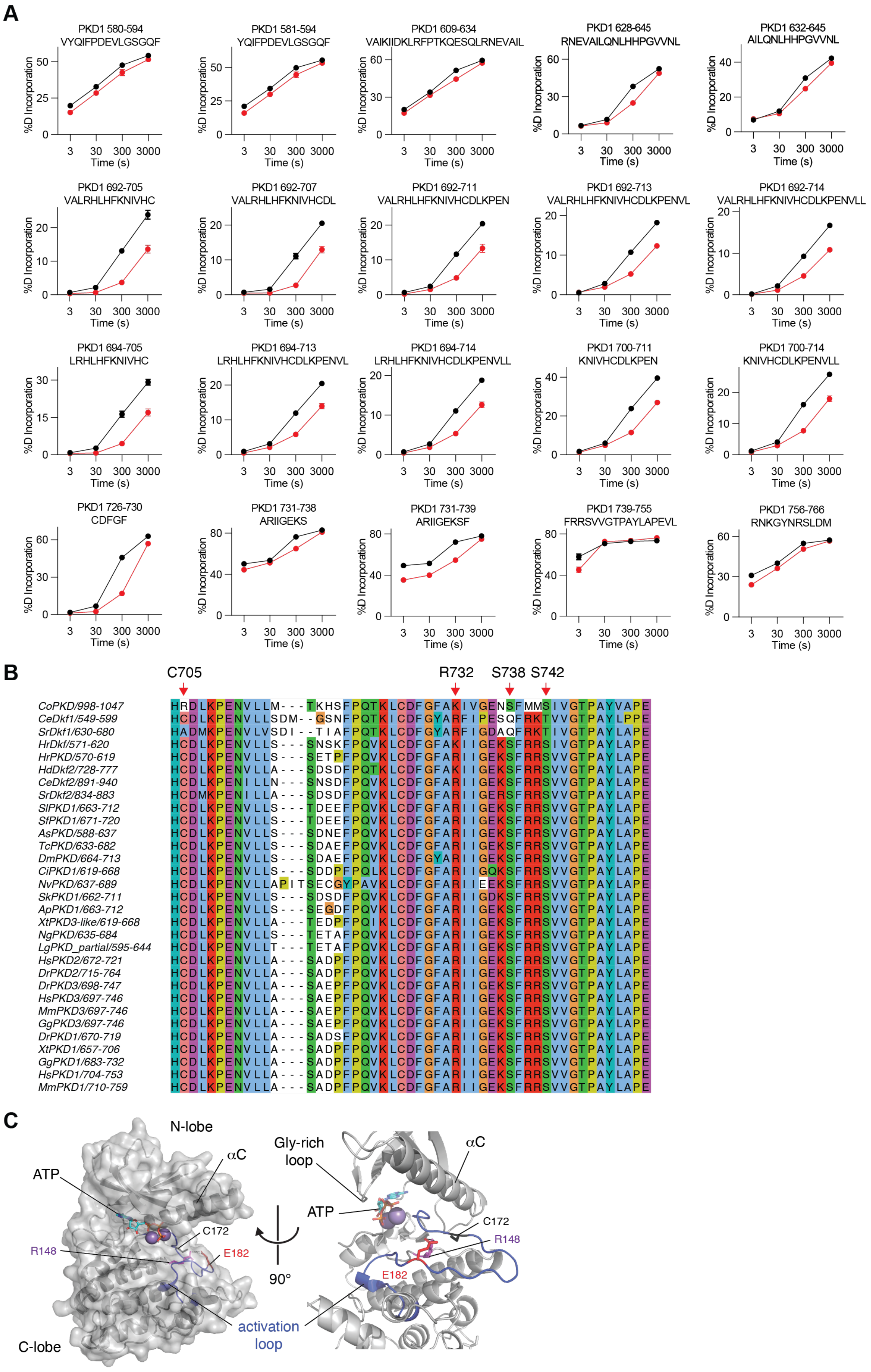

A. Deuterium incorporation for selected PKD1$^{KD}$ peptides. Error bars are the standard deviation of 3 biologically independent experiments.
B. Sequence conservation in the catalytic and activation loops of PKD orthologs. Key residues in activation loop phosphate coordination are indicated with red arrows.
C. Coordination of the activation loop (blue cartoon) in phosphorylase kinase. E182 (red sticks) occupies the position of the canonical phosphoserine in PKD1 and is coordinated by R148 (purple sticks) in the HRD motif.

**Figure S6. Activation loop autophosphorylation increases PKD1 catalytic activity.**

**A**

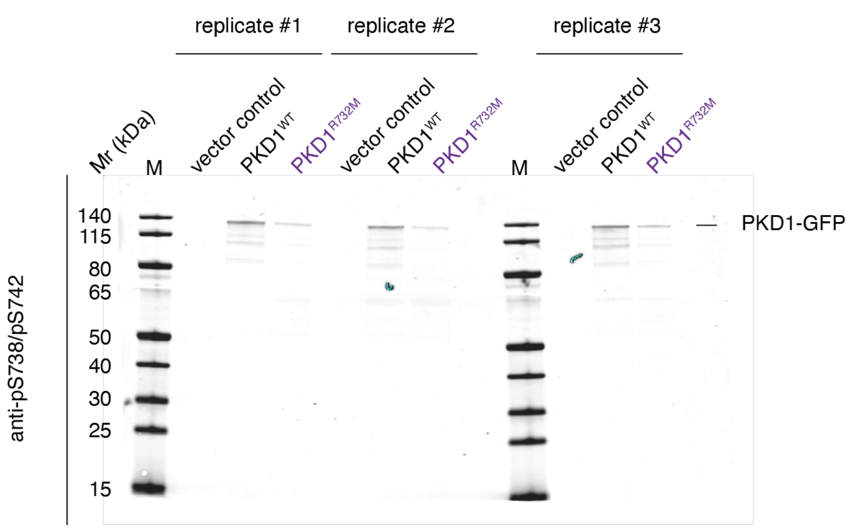


**B**

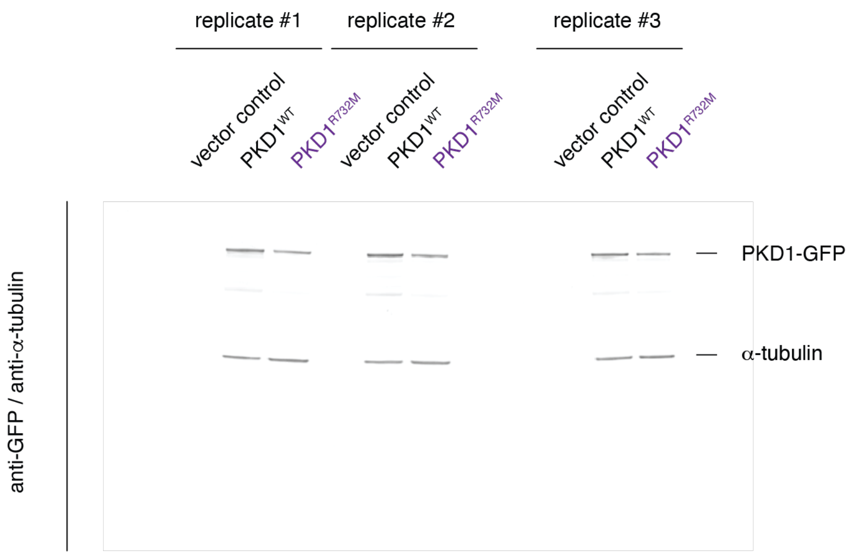


A. Western blots quantified in Figure 3F. Blot probed with anti-PKD1 pS738/pS742 antibody.

B. Western blots quantified in Figure 3F. Blots probed with anti-GFP and anti-α-tubulin antibodies.

**Figure S7. PKD auto- and substrate phosphorylation are mechanistically distinct.**

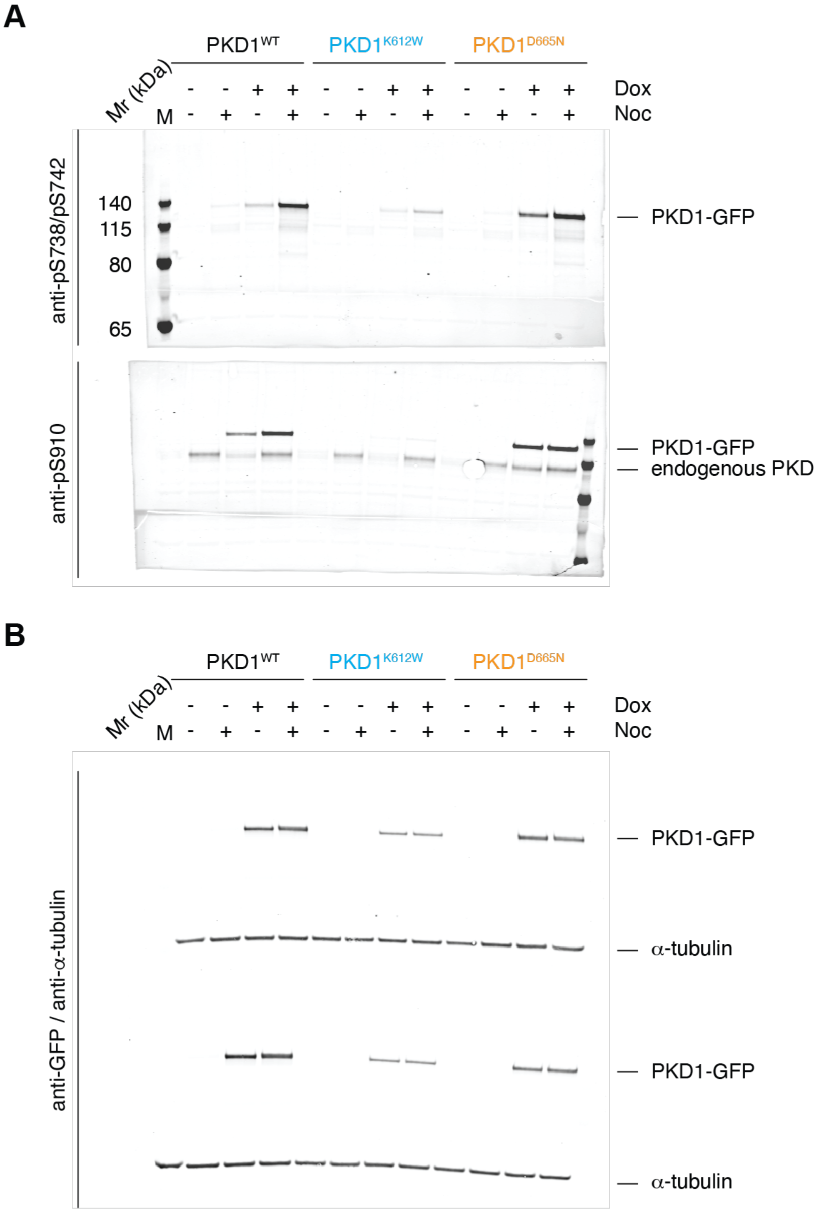


A. Western blots quantified in Figure 4E-F. Upper blot probed with anti-PKD1 pS738/pS742 antibody. Lower blot probed with anti-PKD1 pS910 antibody.

B. Western blots quantified in Figure 4E-F. Blot probed with anti-GFP and anti-α-tubulin antibodies.

**Figure S8. Substrate binding-deficient and constitutively dimeric PKD1 block secretion.**

A

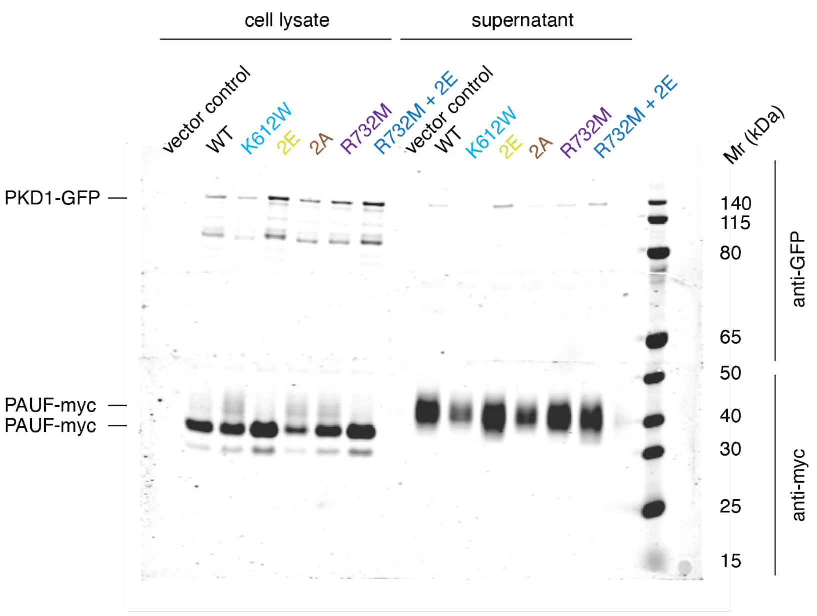


B

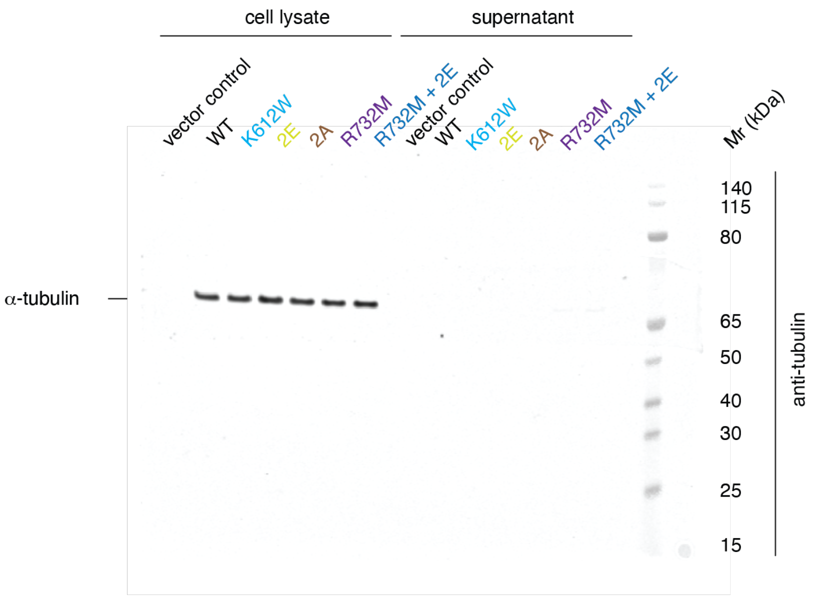


A. Western blot quantified in Figure 6C. Upper blot probed with anti-GFP antibody. Lower blot probed with anti-myc antibody.

B. Western blot loading control for Figure 6C. Blot probed with anti-α-tubulin antibody.

**Figure S9. Autoregulation of membrane binding depends on ATP, but not dimerization.**

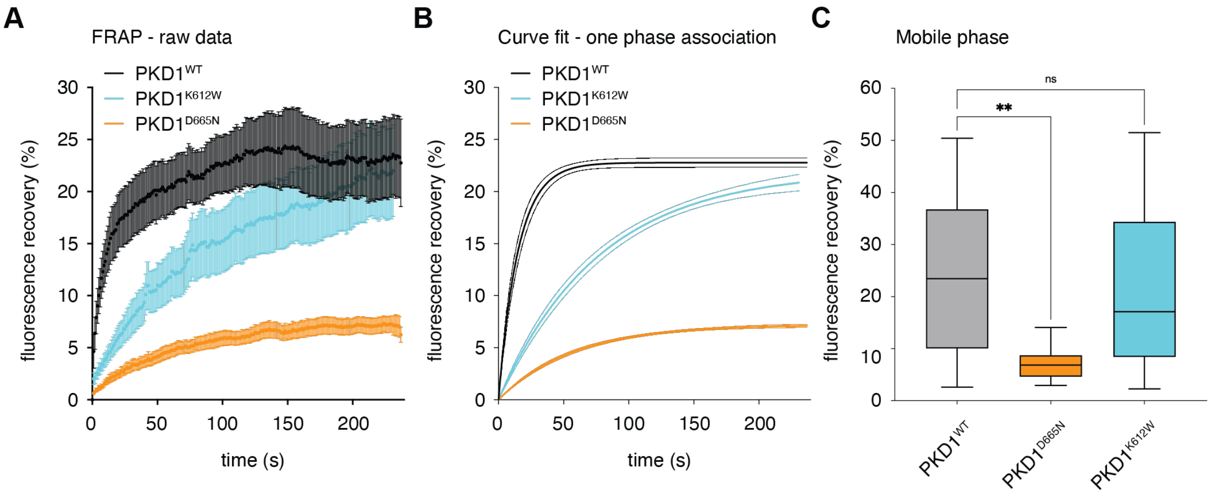


A. Raw FRAP data for PKD1$^{WT}$ (black), PKD1$^{K612W}$ (cyan), and PKD1$^{D665N}$ (orange) obtained from stable Flp-In TRex HeLa PKD1-GFP cells.

B. One-phase association curve fits to the FRAP data in panel A.

C. Mobile phase (maximum fluorescence recovery) was determined by fitting the curves shown in A to a one-phase exponential equation (B). The box plot shows the results of two independent experiments. Center lines show the medians; box limits indicate the25th and 75th percentiles as determined by GraphPad Prism 9 software; whiskers extend 1.5 times the interquartile range from the 25th and 75$^{th}$ percentiles, outliers are represented by dots. n = 14, 16, 13 sample points. The significance of differences was analyzed by an ordinary one-way ANOVA followed by Sidak's multiple comparison test. **$p<0.01$.

**Supplementary Table S1. EDC cross-linked peptides (PDK1$^{KD}$).**

| Crosslinked residues | | Crosslinked peptides | | Abundance | | |
|---|---|---|---|---|---|---|
| | | | | Mono-mer | Di-mer | Total |
| 736 | 622 | IIGEK(4) | FPTKQESQLR(4) | 7 | 3 | 10 |
| 615 | 622 | IIDK(3) | FPTKQESQLR(4) | 1 | 2* | 3 |
| 736 | 758 | IIGEK(4) | NKGYNR(2) | 6 | 2 | 8 |
| 737 | 668 | IIGEKSFR(5) | LHGDMLEMILSSEK(7) | 4 | 0 | 4 |
| 616 | 624 | IIDKLR(4) | QESQLR(2) | 5 | 5* | 10 |
| 737 | 624 | IIGEKSFR(5) | QESQLR(2) | 1 | 0 | 1 |
| 736 | 737 | IIGEKSFRR(4)(5) | | 6 | 0 | 6 |
| 708 | 710 | NIVHCDLKPENVLLASADPFPQVK(8)(10) | | 7 | 2 | 9 |
| 852 | 854 | ELECKIGER(3)(5) | | 2 | 0 | 2 |
| 622 | 624 | FPTKQESQLR(4)(6) | | 18 | 0 | 18 |
| 674 | 675 | LHGDMLEMILSSEKGR(13)(14) | | 1 | 0 | 1 |
| 612 | 615 | DVAIKIIDK(5)(8) | | 1 | 0 | 1 |
| 850 | 854 | ELECKIGER(1)(5) | | 2 | 0 | 2 |
| 831 | 622 | YSVDK(4) | FPTKQESQLR(4) | 0 | 2 | 2 |
| 668 | 622 | LHGDMLEMILSSEK(7) | FPTKQESQLR(4) | 0 | 4* | 4 |
| 708 | 718 | NIVHCDLKPENVLLASADPFPQVK(8)(18) | | 0 | 3 | 3 |
| 706 | 708 | NIVHCDLKPENVLLASADPFPQVK(6)(8) | | 0 | 1 | 1 |

*cross-links used as experimental restraints in Rosetta

**Supplementary Table S2. HDX Data Summary**

| | |
|---|---|
| Protein Data Set | PKD1cat WT/Mutant |
| HDX reaction details | %D2O=71.3%<br>pH(read)= 7.5<br>Temp= 18°C |
| HDX time course | 3s, 30s, 300s, 3000s |
| HDX controls | N/A |
| Back-exchange | Corrected based on %D2O |
| Number of peptides | 128 |
| Sequence coverage | 97.9% |
| Average peptide length / Redundancy | Length = 13.3<br>Redundancy = 4.9 |
| Replicates | 3 |
| Repeatability | Average StDev = 6.0% |
| Significant differences in HDX | >5% and >0.5 Da and unpaired t-test <0.01 |
